# Breaking the rule: An exceptional Y chromosome introgression between deeply divergent primate species

**DOI:** 10.1101/2024.03.07.583898

**Authors:** Axel Jensen, Emma R. Horton, Junior Amboko, Stacy-Anne Parke, John A. Hart, Anthony Tosi, Katerina Guschanski, Kate M. Detwiler

**Affiliations:** Department of Ecology and Genetics, Animal Ecology, Uppsala University, Uppsala SE-75236, Sweden; Department of Biological Sciences, Florida Atlantic University, 777 Glades Road, Boca Raton, FL, 33431, USA; Department of Anthropology, Florida Atlantic University, 777 Glades Road, Boca Raton, FL, 33431, USA; Department of Anthropology, New York University, 25 Waverly Place, New York, 10003, New York, USA; New York Consortium in Evolutionary Primatology, New York, 10024, New York, USA; Lukuru Wildlife Research Foundation, 1235 Avenue des Poids Lourds, Kinshasa, Democratic Republic of Congo; Department of Anthropology and School of Biomedical Sciences, Kent State University, Kent, Ohio 44242, USA; School of Biological Sciences, Institute of Ecology and Evolution, University of Edinburgh, Edinburgh, UK

**Author notes:** These authors contributed equally to this work (Katerina Guschanski, Kate M. Detwiler).

## Abstract

Hybridization between distinct lineages is widespread in nature, with important implications for adaptation and speciation. If hybrids are fertile, alleles can introgress across species boundaries. Hybrid fitness is often sex dependent, with heterogametic hybrids showing lower fitness than homogametic individuals, a phenomenon known as Haldane’s rule. Consequently, loci inherited strictly through the heterogametic sex rarely introgress. We focus on the Y-chromosomal history of guenons (tribe Cercopithecini), a group of African primates that hybridized extensively throughout their evolutionary history. Although our inferences suggest that Haldane’s rule is generally in place, we uncover a remarkable Y chromosome introgression event from *Cercopithecus mitis* into *C. denti*, which occurred ca. six million years after their initial divergence. Only a weak signal of autosomal gene flow is present, suggesting that the introgressing Y chromosome reached fixation in *C. denti* from a very low initial frequency. We use simulations to explore the evolutionary mechanisms at play, and find that selection is needed to achieve fixation of the novel Y chromosome. We identify a number of fixed protein differences between the introgressed and ancestral Y, with meiotic drive being an alternative mechanism to adaptive introgression. Our results show that the ability to produce viable and fertile heterogametic hybrids persisted for six million years in guenons, providing a remarkable exception to Haldane’s rule.

## Introduction

Interspecific hybridization is widespread in nature, and is an increasingly acknowledged evolutionary force (Abbott et al., 2013). However, when divergent lineages interbreed, the resulting hybrids often show a sex-specific reduction in fitness due to genomic incompatibilities. This phenomenon was first described a century ago by Haldane, who stated that ‘when in the offspring of two different animal races one sex is absent, rare, or sterile, that sex is the heterozygous sex’ (Haldane, 1922). Haldane’s rule has been confirmed by observations in nature and in laboratory crosses across diverse species with sex chromosomes, including animals and plants (Brothers and Delph, 2010; Ottenburghs, 2022; Schilthuizen et al., 2011). As a consequence, gene flow between divergent lineages is more likely to occur through homogametic hybrid individuals, and loci that are strictly inherited through the heterogametic sex are unlikely to introgress (Cortés-Ortiz et al., 2019; Geraldes et al., 2008; Vanlerberghe et al., 1986). Consequently, hybridization frequently leads to phylogenetic discordance between autosomes and loci inherited only through the homogametic sex (like the mitochondrial genome [mtDNA] in mammals; (Bastos-Silveira et al., 2012; Good et al., 2015; Jensen et al., 2023)), whereas the phylogeny of loci limited to the heterogametic sex (e.g. Y/W chromosomes) tend to agree with the autosomal tree (Cortés-Ortiz et al., 2019; Geraldes et al., 2008).

Among several explanations for Haldane’s rule, the dominance theory is most frequently invoked (Schilthuizen et al., 2011). It states that any incompatibilities involving the sex chromosomes will be exposed to selection in heterogametic hybrid individuals, whereas homogametic hybrids only suffer the fitness cost of dominant incompatible alleles. As genomic incompatibilities accumulate with divergence time, isolated lineages eventually reach a “point of no return”, beyond which these incompatibilities cannot be overcome. At this stage, the fitness of hybrid heterogametic individuals is zero, and the transmission of the interspecific Y/W chromosomes stops. This certainly seems to hold in mammals, for which only a few cases of Y chromosome introgression have been reported (Chiou et al., 2021; Macholán et al., 2019; Tosi et al., 2002; Wheeldon et al., 2013), all between relatively close lineages (< 3 million years divergent).

Here, we investigate the presence of Y chromosome introgression among lineages that experienced ample gene flow throughout their evolutionary history. Specifically, we focus on guenons, a group of African primates with more than 30 recognized species (Grubb et al., 2003; IUCN, 2022; Lo Bianco et al., 2017). Guenons are renowned for their ability to hybridize (Detwiler, 2019; Detwiler et al., 2005; Jong and Butynski, 2010), and interspecific gene flow was highly prevalent in their past (Ayoola et al., 2021; Jensen et al., 2023; van der Valk et al., 2020). Furthermore, male guenons typically disperse upon maturation, whereas females remain resident (Isbell and Enstam, 2007), which increases the chance for the Y chromosome introgression across species boundaries. All this makes guenons an ideal system to study the impact of Haldane’s rule along a speciation continuum. Although we find that Y chromosomal and autosomal phylogenies generally agree in guenons, in line with the expectations under Haldane’s rule, we uncovered a Y chromosome introgression event between deeply divergent lineages, at a temporal scale that is unprecedented in mammals. Using simulations, we demonstrate that the Y chromosome was introgressed at a low initial frequency, and driven to fixation by positive selection.

## Results

### Sequencing and Genotyping

We expanded a recently published dataset (Jensen et al., 2023; Kuderna et al., 2023) by generating whole genome sequencing data from two previously unsequenced guenon species: *Cercopithecus denti* (one male, one female) and *C. wolfi* (two males, one female). We also sequenced one male each of *C. mitis* (ssp. *stuhlmanni*) and *C. hamlyni*. Together with other published guenon genomes (Ayoola et al., 2021; van der Valk et al., 2020) and two outgroup species from the sister tribe Papionini (Kuderna et al., 2023), our final data set contained 57 samples from 26 species (Table S1). All genomes were sequenced to high coverage (≥19 x, Table S1) using the Illumina platform. The reads were mapped and genotyped against the rhesus macaque (*Macaca mulatta*) reference genome (Mmul_10, GenBank: GCA_014858485.1), yielding on average 2.2 genotyped gigabases (Gb) per sample.

### Phylogenies from markers of different inheritance modes reveal a deep autosomal vs. Y-chromosomal conflict in *Cercopithecus denti*

We inferred the guenon species tree with ASTRAL (Zhang et al., 2018) from 5,037 independent, autosomal gene trees (Figure 1A). The species tree topology was in agreement with that reported by (Jensen et al., 2023), and all genera, species groups, and species, where more than a single sample was sequenced, were monophyletic. As expected, the newly sequenced *Cercopithecus denti* and *C. wolfi* show a sister species relationship and cluster together with the phenotypically similar and geographically close *C. pogonias* (Grubb et al., 2003; IUCN, 2022) hereafter referred to as the eastern *mona* clade (Figure 1A). We also estimated the mitochondrial phylogeny using IQTree (Minh et al., 2020). While *C. denti* and *C. wolfi* showed similar placements as in the species tree, the mitochondrial topology confirmed the previously described, extensive mito-nuclear discordance among guenons (Figure S1), which are the consequence of rampant ancestral gene flow combined with incomplete lineage sorting (ILS) (Jensen et al., 2023).

**Figure 1.**
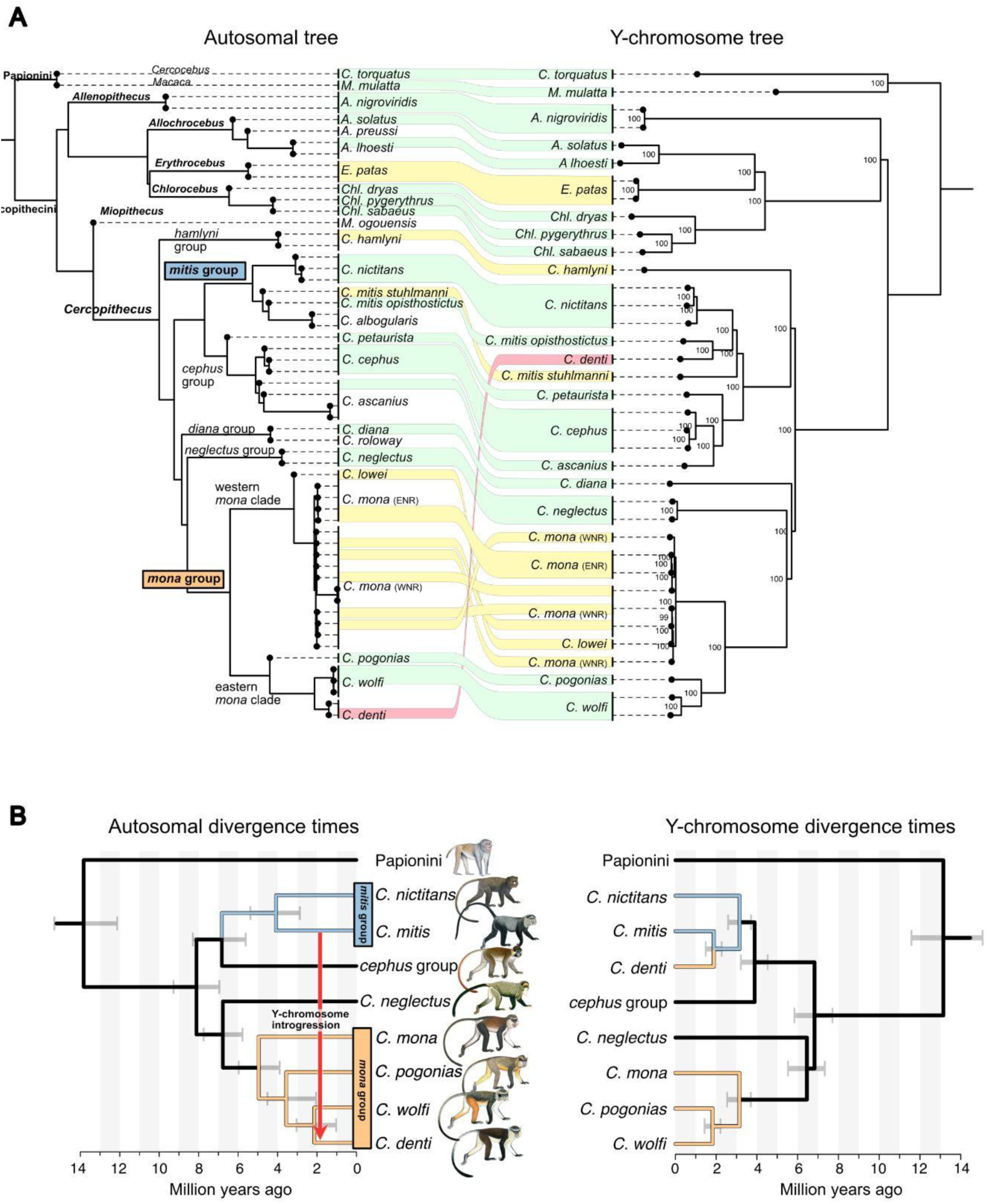
Autosomal and Y-chromosomal phylogenies of guenons. A) Astral coalescence tree constructed from autosomal data (left) and a maximum likelihood tree based on a Y chromosome alignment (right). Connectors show the Y-chromosomal position of male samples relative to the autosomal tree: Concordant phylogenetic positions between the autosomal and the Y-chromosomal tree are shown by green connections. Shallow discordances that can be explained by either ILS or introgression are shown in yellow, whereas the deep discordance for the placement of *C. denti* that cannot be explained by ILS is shown in red. Following Ayoola et al. (2021), we separate *C. mona* in two populations, as indicated by labels in parentheses (ENR and WNR, east/west of the Niger river, respectively). B) MCMCTree divergence date estimates among focal lineages, based on autosomal (left) and Y-chromosomal (right) data. The arrow in the autosomal tree depicts the inferred Y chromosome introgression. Guenon illustrations by Stephen Nash, used with permission.

To infer the Y-chromosomal phylogeny, we constructed a maximum likelihood tree based on a Y chromosome alignment of all males in our data set, using IQTree (Figure 1A, hereafter referred to as the Y-tree). In contrast to the mitochondrial phylogeny, the Y-tree was very similar to the species tree, as would be expected under Haldane’s rule. *Cercopithecus denti*, however, stands out: In stark contrast to its position in the species tree as a *mona* group member, *C. denti* is nested within the *mitis* group on the Y-tree. The closest Y chromosome relative to *C. denti* is *C. mitis opisthostictus* (Figure 1A), with a current distribution range south of *C. denti* (Figure 2). The *mitis* group contains the taxonomically poorly resolved *C. mitis* and *C. albogularis* species, which form a paraphyletic clade in our analyses (Figure 1A). For simplicity, we treat this clade as a single lineage, referred to as *C. mitis* hereafter, and use *C. m. opisthostictus* as the representative of this lineage in downstream analyses. Using MCMCTree (Yang, 2007), we estimated the autosomal divergence time between the *mitis* and *mona* groups to ca. 8 million years ago (Mya), and the Y-chromosomal divergence between *C. denti* and *C. mitis opisthostictus* to ca. 1.9 Mya (Figure 1B, Figures S2-S4). If introgression indeed caused this deep discordance between the species tree and the Y-tree, the Y chromosome must have introgressed between lineages that diverged more than 6 million years earlier.

**Figure 2.**
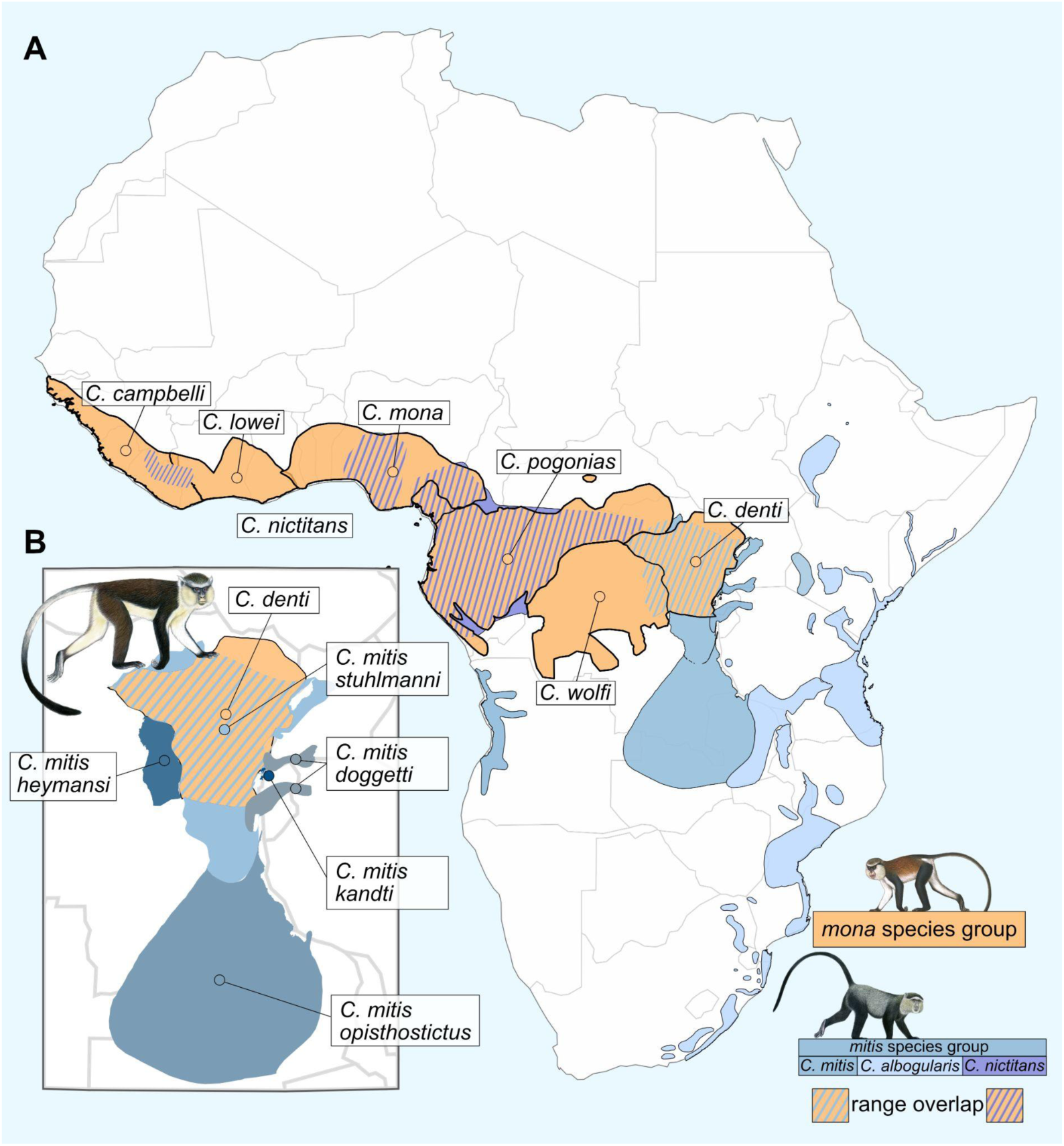
Distribution ranges of the *mona* and *mitis* group lineages. A) Distribution ranges of recognized *mona* and *mitis* group species. B) Zoom-in around the distribution range of *C. denti*, showing its overlap with, and close proximity to, multiple *C. mitis* subspecies. Guenon illustrations by Stephen Nash, used with permission.

Additional discordances between the species tree and the Y-tree were shallow, involving lineages that differ in position across a single speciation node (Figure 1A). Although they may be the result of Y chromosome introgression, such patterns can be expected under ILS alone (Jensen et al., 2023).

### Introgressive hybridization led to the fixation of a divergent Y chromosome

We confirmed that the *mitis*-like Y chromosome is fixed in *C. denti* by sequencing the Y-linked *TSPY* gene in four additional *C. denti* males, sampled across the species’ distribution range (Figure S5, Table S2). To test if the deep discordance between the Y-tree and the species tree was indeed caused by introgression rather than ILS, we compared pairwise nucleotide divergence (*d_XY_*) among *mona* and *mitis* group taxa on the autosomes and the Y chromosome, since introgression and ILS would generate distinct *d_XY_* patterns. For ILS to generate the discordant Y-tree placement of *C. denti*, the Y chromosome must have remained polymorphic and unsorted along the ancestral *mona* and *mitis*/*cephus* group branches (Figure 3A). In this scenario, the Y chromosome of *C. denti, C. mitis, C. nictitans* and the *cephus* group would all coalesce in their common ancestor, producing a greater (or similar) divergence on the Y chromosome compared to the autosomes. Under introgression, the divergence between *C. mitis* and *C. denti* should be much lower on the Y chromosome than on the autosomes (Figure 3B), whereas the divergence between *C. denti* and its autosomal sister *C. wolfi* should be higher on the Y chromosome than on the autosomes. For taxa comparisons not involving *C. denti*, similar Y-chromosomal and autosomal *d_XY_* values are expected.

**Figure 3.**
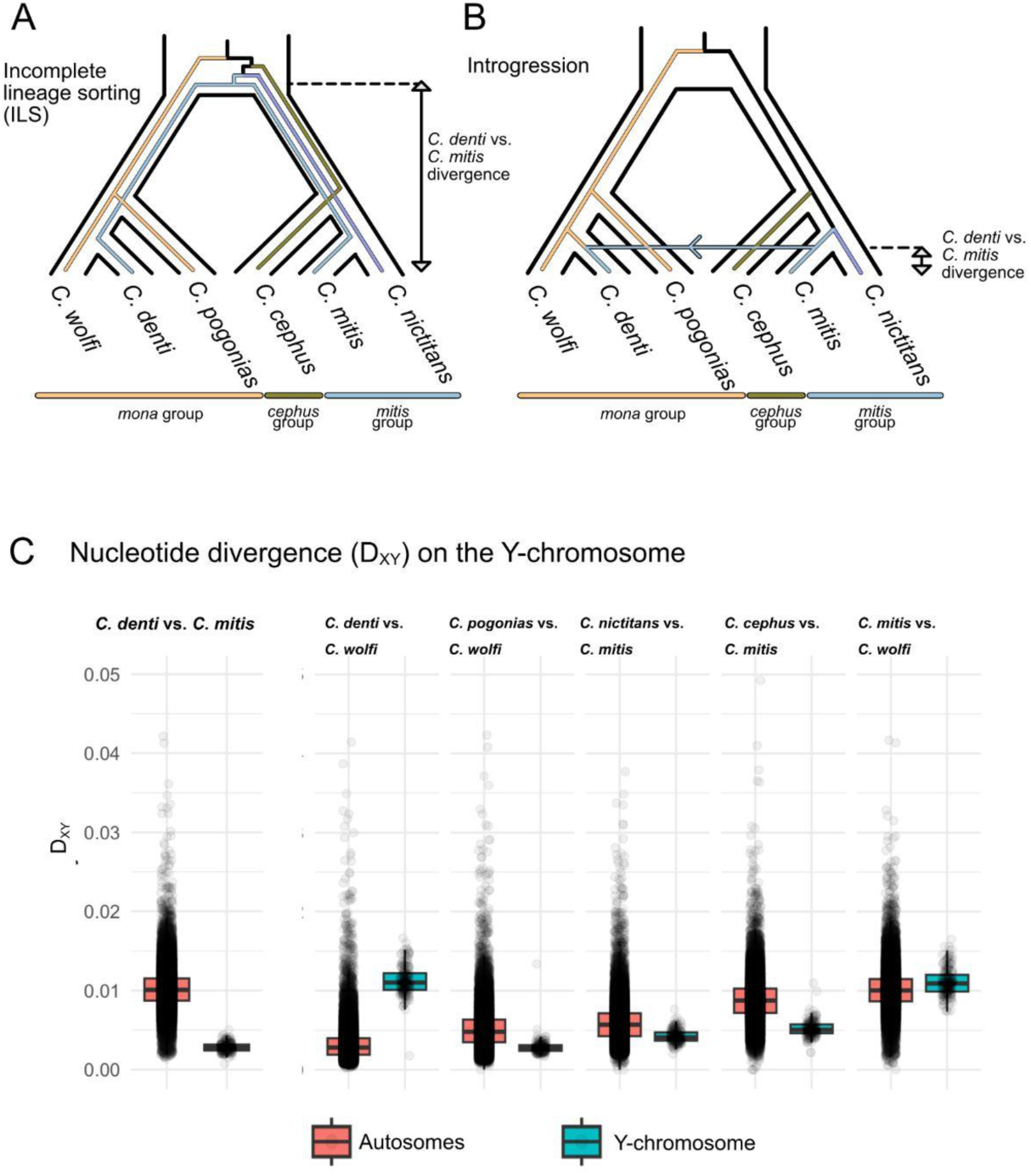
Nucleotide divergence patterns support Y chromosome introgression over ILS in *C. denti*. Schematics of Y chromosome sorting in the *mona* and *mitis*/*cephus* group lineages under ILS **(A)** and introgression **(B)**. Coloured lines represent Y lineages, whereas black outlines show the species tree relationships. Arrow in B corresponds to Y-chromosomal introgression from *C. mitis* into *C. denti*. Double-headed arrow reflects expected nucleotide divergence (d*xy*) for the Y chromosome between *C. denti* and *C. mitis*. **C)** Values of nucleotide divergence (d*xy*) across taxa present in A) and B), supporting introgression of the Y chromosome from *C. mitis* into *C. denti*.

Nucleotide divergence between *C. mitis* and *C. denti* was on average 3.6 times greater on autosomal compared to Y-chromosomal loci, consistent with introgression of the Y chromosome from *C. mitis* into *C. denti* (Figure 3B). Furthermore, the *d_XY_* estimates among *mitis* and *cephus* group species showed similar values for the Y chromosome and the autosomes, suggesting that sorting of the Y chromosome along the ancestral branches of these lineages was complete, hence opposing the expected ILS pattern.

### Low levels of autosomal allele sharing suggests that the Y chromosome introgressed at a low initial frequency

To test for autosomal gene flow corresponding to the Y chromosome introgression from *C. mitis* into *C. denti*, we estimated D-statistics (Durand et al., 2011) for different combinations of *mona* and *mitis* group taxa (Figure 4). Using the rhesus macaque as outgroup, we tested for excess allele sharing (indicative of gene flow) between *C. mitis*/*nictitans* and *C. denti*, compared to *C. mona, C. pogonias* and *C. wolfi*, iterating through all combinations of samples of the respective species. We detected a strong excess of allele sharing between all *mitis* group taxa and *C. denti* compared to *C. mona* (Figure 4A, Table S3). This is consistent with the previously reported allele sharing between the *mitis* group and the eastern *mona* clade (Jensen et al., 2023). However, Jensen et al. (2023) showed that this signal was mainly driven by shared ancestry between the *mitis* and *cephus* groups, as it disappeared in *mitis* but not in *cephus* when considering only variants private to either lineage. Thus, since this gene flow event occurred from *C. cephus* rather than from *C. mitis*, and into the common ancestor of *C. denti*, *C. wolfi* and *C. pogonias*, it is unlikely to have introduced a *mitis*-like Y chromosome only into *C. denti*.

**Figure 4.**
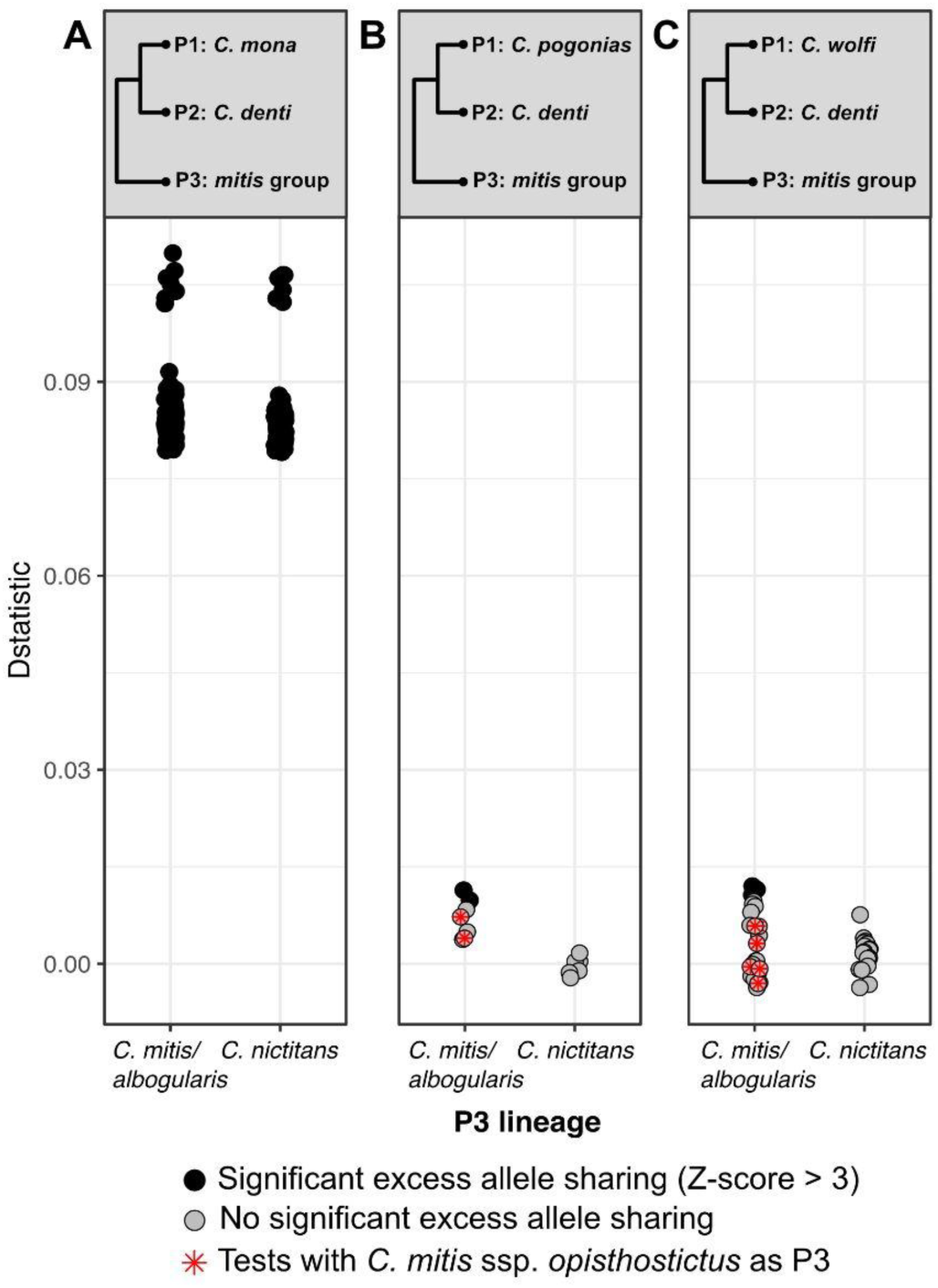
Estimated excess allele sharing between *C. denti* and *mitis* group taxa relative to *C. mona* (A), *C. pogonias* (B), and *C. wolfi* (C). Filled circles show significant D-statistics (Z-score > 3), and points with red asterisks show tests where the *C. m. opisthostictus* was used as P3.

Relative to its closest sister species (*C. pogonias* and *C. wolfi*), we found only weak signals of excess allele sharing between *C. denti* and *C. mitis*, with only five out of 56 tests producing significant D-values (Z-score > 3, Figure 4, Table S3). Furthermore, we found no excess allele sharing between *C. denti* and specifically its closest Y-chromosomal relative *C. m. opisthostictus,* in comparison to *C. wolfi* or *C. pogonias*. Overall, these analyses provide weak support for autosomal gene flow from *C. mitis* into *C. denti* after the split from *C. pogonias* and *C. wolfi*. This suggests that the effective migration rate was low, and the *mitis*-like Y chromosome that is now fixed in *C. denti* was likely introduced at a very low initial frequency.

To identify other genomic regions that may have introgressed from *C. mitis* into *C. denti*, we calculated F_D_ and *d*_XY_ in sliding 10 Kb windows along the autosomes and X-chromosome (Methods). We identified 136 putatively introgressed regions (Figure S6), with the longest being 40 kb in length. These regions overlapped 55 protein coding genes (Table S4), which were not enriched for any gene ontology.

### Genetic Drift is an unlikely driver of the Y chromosome fixation

After establishing that the mitis-like Y chromosome must have introgressed at low initial frequency, we used simulations to investigate if its fixation in *C. denti* requires selection or if it could be explained by drift alone. As a first step, we explored plausible proportions of migration from *C. mitis* into *C. denti* given our empirical D-statistic estimates. We used the multi-species-coalescence-with-introgression (MSci) model implemented in BPP (Flouri et al., 2020) to infer the demographic history of the focal lineages as a basis for subsequent simulations.

We included three gene flow events in our model (Figure 5): (1) from the *mitis* group ancestor into the ancestor of *C. denti*, *C. pogonias* and *C. denti*, (2) from the *C. cephus* lineage to the ancestor of *C. pogonias*, *C denti* and *C. wolfi* (as inferred in (Jensen et al., 2023)), and (3) from *C. mitis* into *C. denti* (corresponding to the Y chromosome introgression). Although there is no direct support for the gene flow event 1 (Jensen et al., 2023), it is possible that it could partially be masked by extensive gene flow from the *cephus* group into the *mona* group (event 2). We therefore conservatively included this event since it could be a source of the Y chromosome introgression (together with ILS) into *C. denti*, and have an effect on the observed D-statistics.

**Figure 5.**
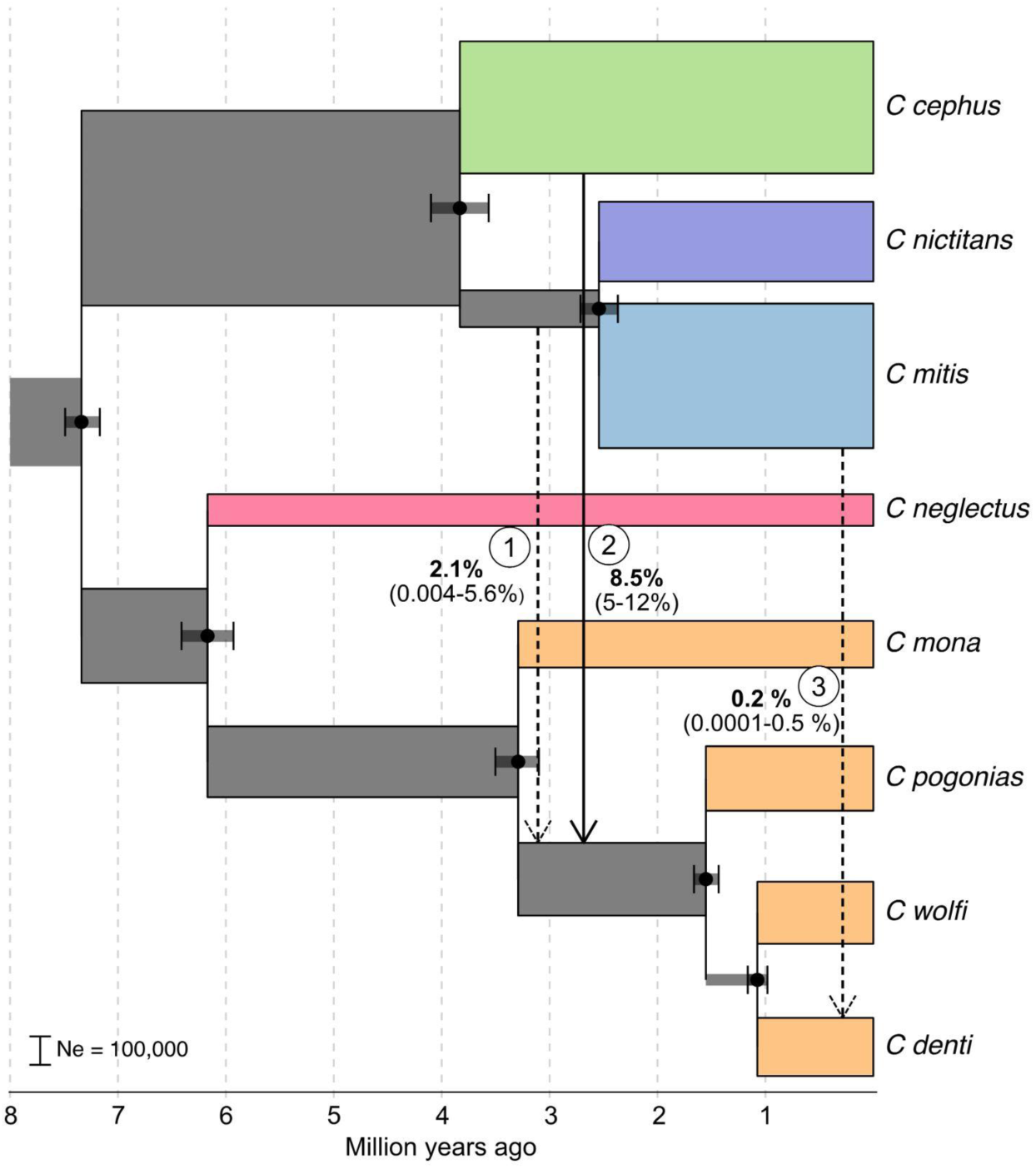
Demographic history of focal lineages as estimated with BPP-MSci. Gene flow events included in the demographic model are shown as arrows, signifying their directionality. Labels next to the arrows show the obtained means and 95 % highest posterior density intervals of estimated migration rate for each event. Dashed arrows indicate gene flow events with low support from D-statistics estimates (Figure 4, (Jensen et al., 2023)). Widths of the branches correspond to inferred effective population sizes (Ne), as indicated by the scale bar.

Two independent runs of BPP-MSci with the same parameters converged on highly similar estimates (Figure S7). Scaling the divergence times using a generation time of 10 years and a mutation rate of 4.82e-9 (Kuderna et al., 2023), resulted in estimates similar to the MCMCTree analysis (Figure 5, Figure 1B), confirming that ca. 6 million years of divergence preceded the Y chromosome introgression from *C. mitis* into *C. denti*.

Ancestral effective population size estimates were generally large (> 100,000, Figure 5), in line with the reported high genetic diversity in guenons (Jensen et al., 2023; Kuderna et al., 2023). The most pronounced gene flow event was, as expected, from *C. cephus* into the eastern *mona* clade (event 2 in Figure 5, migration rate [phi] = 8.5 %, time ~ 2.7 MYA), followed by event 1 (migration rate ca. 2.1 % at ~ 3.1 MYA.). In line with our D-statistics result, the migration rate from *C. mitis* into *C. denti* (event 3 in Figure 5) was low (phi = 0.2 % ~ 0.3 MYA). Switching the order of the gene flow events 1 and 2 resulted in similar estimates overall, but lower migration proportions in event 1 (Figure S8, phi = 0.8 %).

Using the estimated demographic parameters (Figure 5), we next performed simulations in msprime (Baumdicker et al., 2022) to infer a migration rate from *C. mitis* into *C. denti* that is compatible with our empirical D-statistic estimates. We conservatively included both ancestral gene flow events 1 and 2. For the autosomes, we tested a range of migration proportions from *C. mitis* into *C. denti* (0-1%, with a stepwise increase of 0.05 %) and calculated D-statistics between *C. denti* and *C. mitis* compared to *C. wolfi*. The gene flow was set to occur shortly (10,000 generations) after the split between *C. denti* and *C. wolfi* to allow drift to act on the introgressing loci.

Our simulations showed that effective migration rates of > 0.4 % consistently produce significantly positive D-statistics (Z-score > 3, Figure 6A). This migration proportion also generated consistently greater D values than our highest empirically observed D-statistic for *C. mitis* into *C. denti* compared to *C. wolfi*. For computational reasons, the simulated genome length was 100 Mb, i.e. considerably smaller than the primate genome (corresponding to only ~3.5%). As the sensitivity of D-statistics increases with the number of loci (Zheng and Janke, 2018), it is therefore highly unlikely that the effective migration rate from *C. mitis* to *C. denti* (that resulted in Y chromosome introgression) exceeded 0.4 %.

**Figure 6.**
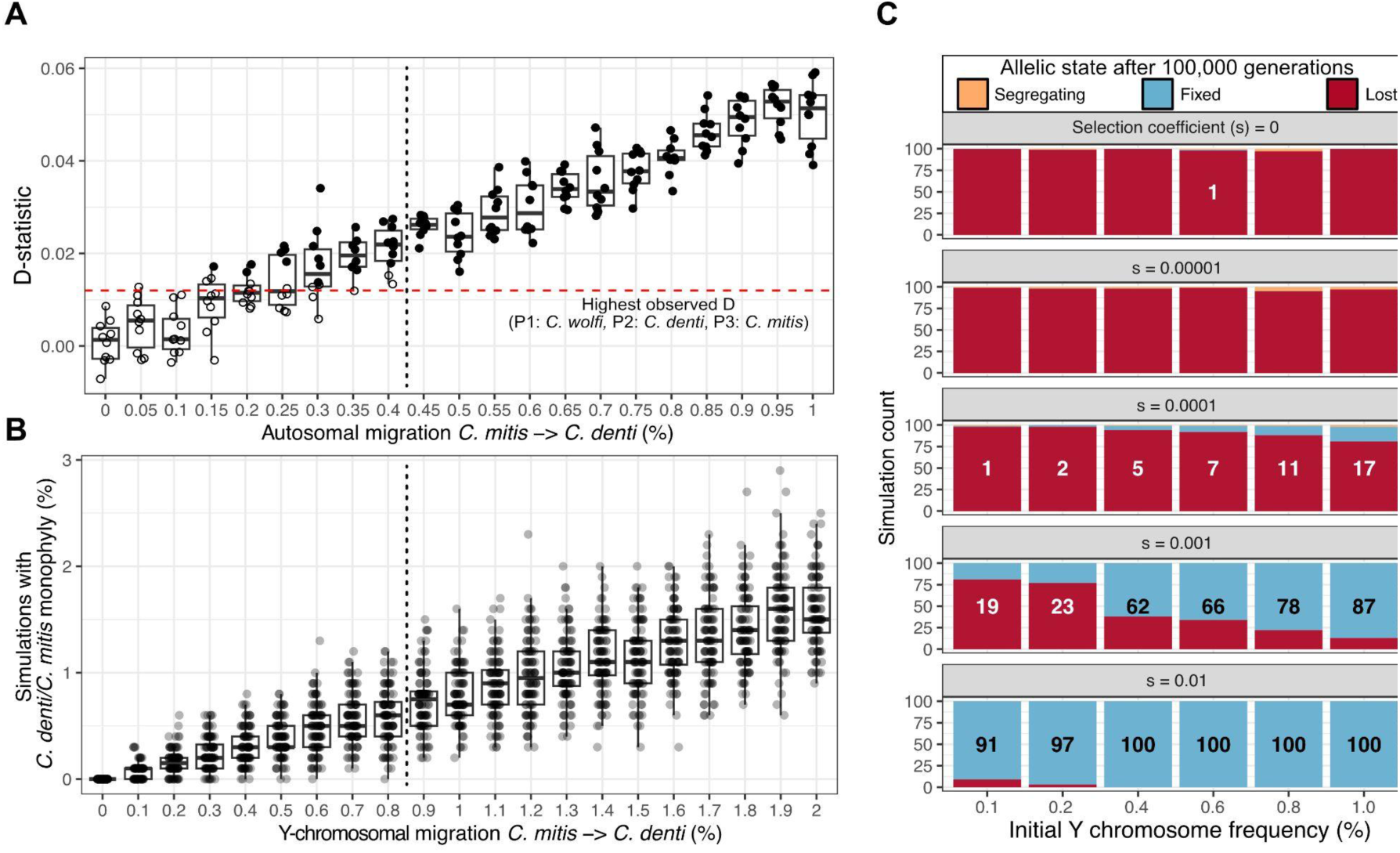
Simulated D-statistics and probability of Y chromosome fixation. **A)** D-statistics (P1=*C. wolfi*, P2=*C. denti*, P3=*C. mitis*, Outgroup=*M. mulatta*) resulting from simulated evolutionary histories with varying amounts of migration from *C. mitis* into *C. denti*. Each circle represents the estimated D-statistics in one simulation, with increasing gene flow proportion along the X-axis. Filled circles represent significantly positive D-statistic tests (Z-score > 3), and the horizontal dashed line depicts the greatest observed D-value from empirical data. Vertical dashed line shows the proportion of gene flow above which all estimates are significant and above the observed D-statistic. **B)** The percentage of simulations (out of 1,000) that resulted in a sister Y-chromosomal relationship between *C. denti* and *C. wolfi* under the same absolute number of migrants as in (A), assuming that all migrants are males. Vertical dashed line is at the same number of migrating individuals as in (A). **C)** The probability of Y chromosome fixation under different initial allele frequencies (equivalent to incoming Y-chromosomal migration, X-axis) and selection coefficients (panels), retrieved using forward simulations in SLiM. Numbers on bars state the count of simulations resulting in fixation of the introgressing Y chromosome, out of a total of 100 replicate simulations.

Next, we estimated the probability of a sister relationship between *C. denti* and *C. mitis* on the Y chromosome, when considering different rates of migration between these two lineages (event 3 in Figure 5). The Y chromosome was simulated as a single, non-recombining locus with effective population sizes of ¼ of the autosomes. We conservatively assumed that all *C. mitis* migrants entering the *C. denti* population were males, leading to an effective Y-chromosomal migration rate twice as high as the autosomal migration rate (0-2%, with a stepwise increase of 0.1 %, to retrieve migration proportions directly equivalent to the autosomal migration rates). We then calculated the proportion of simulations in which *C. denti* and *C. mitis* formed a monophyletic Y-chromosomal clade when randomly sampling one individual per species, which reflects the chance of a sister relationship among these species. We ran 100 replicates, comprising 1000 simulations each, for every migration rate.

Under strictly male mediated gene flow, an autosomal migration rate of 0.4 % corresponds to a Y-chromosomal migration rate of 0.8 % (Figure 6B). At this migration rate, the chance of a sister relationship between *C. denti* and *C. mitis* on the Y chromosome was ~0.6 %, according to our simulations. Even with gene flow proportions that by far exceed the empirical values, neutral drift alone is highly unlikely to drive the introduced Y chromosome to fixation, and a monophyletic Y-chromosomal relationship between *C. denti* and *C. mitis* was rare throughout the tested range of migration (average ≤ 1.6 %). Furthermore, these simulations confirm that ILS alone is extremely unlikely to have caused the discordant Y-chromosomal tree position of *C. denti*: Without migration (Y-chromosomal gene flow of 0 %, Figure 6B), *C. denti* and *C. mitis* never formed a monophyletic clade (out of a total of 100,000 simulations).

### Moderate positive selection is capable of driving the Y chromosome fixation in *C. denti*

To test the ability of selection to drive the introgressed Y chromosome to fixation, we ran forward simulations in the SLiM framework (Haller and Messer, 2019). We simulated a population of 200,000 individuals for 100,000 generations (mimicking the *C. denti* lineage based on our BPP analysis, Figure 5) and traced the frequency of a novel Y-chromosomal allele introduced at different initial frequencies (i.e. introgression proportions), with varying selection coefficients (s).

In line with the coalescent simulations in msprime, the novel Y chromosome was highly unlikely to fix without selection: Drift alone drove the novel allele to fixation a single time across 600 simulations (with initial Y chromosome frequencies ranging between 0.1 and 1 %), and it remained segregating after 100,000 generations in four additional replicates (Figure 6C). However, with a moderate selection coefficient of s = 0.001, the fixation rate of the novel Y chromosome was 62 % already at an initial frequency of 0.4 % (equivalent to the estimated migration rate from BPP-MSci), and 78 % when the initial frequency was 0.8 % (equivalent to the upper bound of migration based on empirical D-statistics and simulations). Hence, these simulations suggest that the introgressing Y chromosome likely had a selective advantage over the ancestral Y, allowing it to reach fixation from a low initial frequency.

### Possible targets of selection on the introgressing Y chromosome

We explored possible targets for positive selection on the introgressing Y chromosome by identifying Y-linked genes with fixed amino acid substitutions in the *mitis* group and *C. denti* compared to *C. pogonias* and *C. wolfi*. Of the 41 annotated Y-chromosomal genes on the macaque reference genome, only 28 contained any genotyped sites in our study genomes, as a consequence of the highly repetitive nature of the Y chromosome (Table S5, Methods). After removing genes with premature stop codons in any of the focal species, we identified 13 genes with at least one fixed amino acid difference in *C. mitis* and *C. denti* compared to *C. wolfi* and *C. pogonias* (Table S5). The highest numbers of substitutions were found in the genes KDM5D and USP9Y, with five and four differences, respectively.

We also searched for evidence of adaptive evolution in these thirteen genes with two different approaches. First, we used codeml implemented in the PAML package (Yang, 2007) to specifically test whether any sites show signature of adaptive evolution along the ancestral *C. mitis*/*C. denti* Y chromosome branch. No such sites were identified (p > 0.25). Second, we used the MEME method implemented in the HyPhy suite (Murrell et al., 2012) to identify sites that show signals of diversifying selection. Four sites were identified as being under diversifying selection (p = 0.080-0.087, Table S6). Three of these, located in DAZA, ZFY and KDM5D, were differentially fixed between the eastern *mona* clade and *C. denti*/*C. mitis*, highlighting these genes as candidate targets for selection.

An alternative to the classical scenario of adaptive introgression is meiotic drive orchestrated by selfish genetic elements. One reported drive mechanism for the Y chromosome is through the expansion of gene copies, which can increase the proportion of Y-bearing sperm relative to X-bearing sperm (Baird et al., 2023; Hughes et al., 2020). This may then be compensated for by copy-number expansions on the X-chromosome, leading to an arms race between the sex chromosomes. During hybridization, the lack of compensatory expansions on the X-chromosome in a naive genome may allow for rapid drive of the introgressing Y chromosome. This scenario was implicated in a Y chromosome introgression between mouse subspecies (Baird et al., 2023; Macholán et al., 2019), and selfish arms-races between the X and Y has been suggested to be a widespread phenomenon in mammals (Hughes et al., 2020). To explore this possibility, we investigated the mapped read coverage in *C. denti* and other guenon species after mapping to the rhesus macaque Y chromosome, as this could indicate a copy number expansion. We identified a region of ca. 1.35 Mb with 2-7 times higher normalized coverage in *C. denti* compared to *C. wolfi* and higher coverage in *C. denti* than in any other *mona* group species (Figure S9-S10). This region contains two genes from the *CDY* gene family (*CDY* and LOC106995435). However, no evidence of higher copy number in this region was present in *C. m. opisthostictus,* the Y-chromosomal sister of *C. denti*, contradicting that this region corresponds to the introgressed drive element. Furthermore, we found no evidence of compensatory copy-number expansions on the *C. denti* X-chromosome relative to *C. wolfi* (Figure S11). The signature of a copy-number expansion in this 1.35 Mb Y-chromosomal region was also present in other guenon species (Figure S12). While it is possible that this or some other undetected Y chromosome region has been implicated in meiotic drive in *C. denti*, our results are inconclusive. Resolving the ampliconic structure of the Y chromosome, which is possible only through the use of long-read sequencing data, will provide a better understanding of these processes (Makova et al., 2023).

## Discussion

In this study, we investigate the prevalence of Y chromosome introgression in a species group that experienced extensive hybridization throughout their evolutionary history. Although the Y-chromosomal phylogeny was generally in agreement with the species tree, we uncover the most distant Y chromosome introgression reported in mammals to date. We explore the evolutionary processes behind this exceptional event that occurred between primate lineages that diverged from each other more than six million years prior to the introgression (corresponding to ca. 600,000 generations of independent evolution). As illustrated by the near-absence of autosomal introgression, the Y chromosome swept to fixation in *C. denti* from a very low initial frequency (≤ 0.8 %). We demonstrate that this is highly unlikely under neutral drift alone, and suggest that a selective advantage of the introgressing Y chromosome drove it to fixation.

The evolutionary history of guenons is characterized by rampant ancestral hybridization, involving deeply divergent lineages that experienced >5 million years of independent evolution and differ in chromosome numbers (Jensen et al., 2023). At least five introgression events of the maternally inherited mitochondrial genome (mtDNA) occurred across guenon species (Figure S1) (Ayoola et al., 2021; Jensen et al., 2023; van der Valk et al., 2020), whereas the phylogeny of the paternally inherited Y chromosome is generally consistent with the species tree (Figure 1A). Guenons show strong male-biased dispersal, and the Y chromosome should thus have many more opportunities to introgress than the mitochondrial genome.

Indeed, we observe a number of shallow discordances between the Y-tree and the species tree (Figure 1A). Although ILS may be an important mechanism in generating such discordances, it is likely that some of them are a consequence of male-biased gene flow. For example, the closely related *Cercopithecus lowei* and the two populations of *C. mona* separated by the Niger river (*C. mona* WNR/ENR in Figure 1A; (Ayoola et al., 2021)) form three well-defined clades in both the mitochondrial and autosomal phylogenies (Figure 1A, S1). In the Y-tree they are intermixed, suggesting that male migration continued after initial population divergence. Similarly, *C. hamlyni*, another species involved in a shallow species tree vs. Y-tree discordance (Figure 1A), has experienced gene flow with the ancestor of the *mitis* species group (Jensen et al., 2023). It is possible that *C. hamlyni* received gene flow already from the ancestor of the *mitis* and *cephus* species groups, making introgression a plausible explanation of its Y-tree placement. Finally, *Erythrocebus patas* shows variation in phylogenetic placement using different marker combinations and approaches, likely as a result of rapid speciation and pronounced ILS (Jensen et al., 2023; Kuderna et al., 2023). Even if Y chromosome introgression indeed caused these shallow discrepancies, none of the involved lineages would have been more than ca. 3 million years divergent at the time of introgression (e.g., *C. lowei* vs. *C. mona*). Although Y chromosome introgression is rare already on such time scales, comparable events have been reported previously. For instance, two species of macaques (*Macaca*) experienced Y chromosomal introgression ca. 2-3 million years after they diverged (Matsudaira et al., 2018; Tosi et al., 2002). Y chromosome introgression has also been reported among baboons (*Papio*, (Chiou et al., 2021)), canids (*Canidae*, (Wheeldon et al., 2013)), and mice (*Mus*, (Macholán et al., 2019)), all involving lineages less than 1 million years divergent.

The prevalence of shallow Y-tree vs. species tree discordances confirm that the rampant gene flow experienced by guenons throughout their evolutionary history creates opportunities for Y-chromosomal introgression on a level rarely reported in other taxa. In contrast to the frequent mtDNA transfers, however, introgression events of more distant Y chromosomes were absent, with one notable exception. This strongly suggests that male hybrids are affected by genomic incompatibilities to a greater degree than females. In this context, the Y chromosome introgression into *C. denti* from the deeply divergent *C. mitis* constitutes a remarkable exception to Haldane’s rule. For comparison, (Allen et al., 2020) explored sequence divergence in the mitochondrial cytochrome *b* gene (*CYTB*) as a predictor for Haldane’s rule, and found that fertile heterogametic hybrids were completely absent at divergences higher than 8 %. Based on the contemporary *CYTB* divergence between *C. denti* and *C. mitis* of ~13-14 %, these lineages showed a divergence of >10 % at the time of Y chromosome introgression (assuming a constant substitution rate). For context, this is comparable to the contemporary sequence divergence between humans and chimpanzees (~10.8-11 %, Allen et al. 2020), and further highlights the uniqueness of this introgression event.

Our analyses unambiguously suggest that selection was required to achieve fixation of the introgressing Y chromosome. As potential targets of positive selection, we identified several genes with fixed non-synonymous substitutions between the introgressed and ancestral Y chromosomes. The largest number of amino acid substitutions were found in *KDM5D* and *USP9Y* which, as most Y-encoded genes, have important functions in spermatogenesis (Krausz and Casamonti, 2017; Subrini and Turner, 2021). Mutations or deletions of these genes have been reported to reduce male fertility in humans (Krausz and Casamonti, 2017; Sun et al., 1999). Sperm competition is an important selective force in many primates (Harcourt et al., 1995, 1981; Hirst et al., 2023) and likely also in guenons, where influx of males into the social groups during the breeding season has been reported (Isbell and Enstam, 2007). Therefore, the two detected genes are plausible subjects for adaptive introgression. In line with this, our results also suggest that one of the fixed sites in *KDM5D* evolved under diversifying selection.

Another possible mechanism behind the Y chromosome fixation is meiotic drive. Selfish genetic elements on the sex-chromosomes may act to increase the fertilization success of X or Y-bearing sperm, leading to an arms race between X and Y-linked genes (Baird et al., 2023; Hughes et al., 2020). Meiotic drive was implicated in the asymmetric Y chromosome introgression in a hybrid zone of house mouse (*Mus musculus*) subspecies (Baird et al., 2023), driven by a copy number expansion of the Y-linked *SLY* gene. If not compensated for by copy number increase of the homologous X-linked *SLX* gene, the *SLY* expansion leads to an excess of Y-bearing sperm and a male-biased sex-ratio. Although *SLY/SLX* are specific to the mouse/rat lineage, similar cases of co-amplification of X/Y-linked genes have been described in other mammals (Hughes et al., 2020). We detected a Y-chromosomal region showing signatures of extensive copy number variation among guenons, containing two genes from the *CDY* gene family. A rapid copy-number expansion in *CDY* genes was also reported in orangutan (genus *Pongo*) relative to other great apes (Vegesna et al., 2020). Although the *C. denti* Y chromosome appeared to have higher coverage in this region compared to the other *mona* group lineages, the closest *C. mitis* Y chromosome does not show such a signature, which makes it unlikely that this region constitutes an introgressing drive element. We are currently lacking data on much of the *mitis* species group variation, and it is possible that the actual source of the Y chromosome is an unsampled relative of *C. m. opisthostictus*. If so, it is possible that the copy number expansion occurred in this unrepresented lineage, as copy number expansions in this region were highly variable and species-specific. Therefore, we cannot exclude meiotic drive as the underlying mechanism, and highlight the guenon Y chromosome as an intriguing subject for future studies, preferentially using long-read data.

Whichever mechanism is at play, our estimates of the selection coefficient needed to achieve Y chromosome fixation are likely underestimated. In our simulations, the introgressing Y chromosome is modeled as a neutral locus, whereas the underlying mechanism behind Haldane’s rule results in selection against the foreign Y chromosome. Therefore, the selective advantage of the introgressing Y must have been strong enough to achieve fixation in the presence of negative selection.

Interspecific gene flow between divergent lineages is a well-documented phenomenon (Detwiler, 2019; Figueiró et al., 2017; Jensen et al., 2023; Meier et al., 2017). As a consequence of Haldane’s rule, the backcrossing required for introgression to occur is expected to be mediated mainly through F1 hybrids of the homogametic sex (females in mammals), since the heterogametic hybrids are typically less fit. Hence, Y chromosome introgressions are exceptionally rare in mammals, and if present, occur between closely related lineages that have accumulated few genomic differences. This study thus provides a remarkable exception, where a selectively advantageous Y chromosome introgressed from *Cercopithecus mitis* into *C. denti*, showing that fertile male hybrids were produced between lineages that diverged more than six million years prior. Although the exact mechanisms that facilitated the introgression and enabled the fixation of this Y chromosome remain a mystery at this stage, we propose that selection either on coding genes or drive elements must have been strong enough to overcome the negative selection against the heterogametic hybrid.

## Methods

### Sampling, DNA extraction and sequencing

Non-invasive tissue samples of *C. mitis, C. hamlyni, C. wolfi* and *C. denti* were collected opportunistically from deceased individuals in the Democratic Republic of Congo. We collected an additional non-invasive fecal sample of a male *C. denti* from Nyungwe National Park, Rwanda. Samples were stored in RNAlater or 94% ethanol. Sample collection and export to the United States was carried out under approved country-specific research permits. Total genomic DNA was extracted using the DNeasy Blood & Tissue Kit (Qiagen 69504; Germantown, MD) or QiaAMP DNA Stool Mini Kit (Qiagen 51504; Germantown, MD) following manufacturer protocols. DNA extracts were sent to the Science for Life Laboratory at Uppsala University, Sweden where library preparation and sequencing was performed by the SNP&SEQ Platform using the TruSeq PCRfree DNA library preparation kit (Illumina Inc.). The libraries were sequenced on the NovaSeq 6000 platform, aiming at a sequencing depth of ca. 30 X. In addition, we obtained published medium to high coverage whole genome sequences from 50 additional individuals of 23 species, including two outgroup species from the sister tribe Papionini, adding up to a total of 57 genomes from 26 species (Ayoola et al., 2021; Kuderna et al., 2023; van der Valk et al., 2020)(Table S1).

### Mapping and Variant Calling

We followed the Genome Analysis Toolkit best practices workflow to process the data (Caetano-Anolles, 2023). Briefly, we added read group information and marked adapters with Picard/2.23.4 and aligned the processed reads to the rhesus macaque *Macaca mulatta* reference genome (Mmul_10, GCF_003339765.1) using the mem algorithm in bwa/0.7.17 (Li and Durbin, 2009). The mapped BAM-files were sorted and deduplicated using Picard, and mapping quality and depth assessed with QualiMap/2.2.1 (García-Alcalde et al., 2012). Next, we used GATK/4.2 to first call genotype per samples with HaplotypeCaller in GVCF mode, which were then combined with CombineGVCFs and jointly genotyped with GenotypeGVCFs, set to output also invariant sites. Indels were excluded, and single nucleotide variants were filtered using gatk VariantFiltration following the recommended exclusion criteria (QD < 2.0, QUAL < 30.0, SOR > 3.0, FS > 60.0, MQ < 40.0, MQRankSum < −12.5, ReadPosRankSum < −8.0). Additionally, we used custom python scripts to mask heterozygous sites with minor allele read support < 0.25, and sites with less than half or more than twice the genome-wide average read depth for each sample (for the X and Y, we used the chromosome-wide average when calculating these cutoffs). Repetitive regions were identified and excluded following the SNPable regions pipeline (https://lh3lh3.users.sourceforge.net/snpable.shtml). Last, Y-chromosomal sites where any sample was called as heterozygous were removed.

### Phylogenetic Analyses and Divergence Date Estimates

We used ASTRAL/5.7.4 (Zhang et al., 2018) to infer the autosomal phylogeny for all species under the multispecies coalescent model. ASTRAL takes independent gene trees as input, for which purpose we sampled a 25 Kb alignment every 500 Kb and constructed a maximum likelihood tree with IQTREE/2.2.2.6 (Minh et al., 2020) with 1,000 rapid bootstraps, using the GTR model. To infer the mitochondrial phylogeny, we assembled and annotated the mitochondrial genomes (mtDNA) of all samples with MitoFinder/1.4.1 (Allio et al., 2020), after trimming the raw reads with TRIMMOMATIC/0.39 (Bolger et al., 2014), using a published *Chlorocebus sabaeus* mtDNA as reference (NC_008066.1). Each mtDNA genome was then divided into 42 partitions: 1st, 2nd and 3rd codon positions of 13 protein coding genes, two rRNA and 22 concatenated tRNA, which were individually aligned using MAFFT/7.407 (Katoh et al., 2019). The best partitioning scheme and model was evaluated using the model test implemented in IQTREE, which was then also used to construct a maximum likelihood tree with 1,000 rapid bootstraps. The Y-chromosomal phylogeny was constructed by first converting the genotypes to a chromosome-wide alignment, removing sites with more than 10% missing data, resulting in an alignment of 414,466 bp. A maximum likelihood tree was constructed in IQTREE using the GTR+F model of substitutions, performing 1,000 rapid bootstraps.

We estimated the divergence dates on the autosomes and Y chromosome separately using MCMCTree as implemented in PAML/4.9j (Yang, 2007). We used a single sample per species for these analyses, choosing the sample with the least amount of missing data across the autosomes and Y chromosome, respectively. For the autosomal data, we sampled 10 loci of 5 Kb each, requiring them to be located at least 10 kb from the nearest gene to minimize biases from selection, which were then treated as individual partitions in the analysis. We ran two independent MCMCTree runs, using the correlated rates clock model and sampling every 100 iteration after discarding the first 10,000 as burnin, until a total of 20,000 samples were retrieved. Due to the large heterogeneity expected across the autosomes from, e.g., rate variation or gene flow, we ran 10 replicates (i.e., sampling 10 new loci). The results of each run were first analyzed independently, and subsequently merged and summarized across all runs. Effective sampling sizes (ESS) were obtained using tracer/1.7.1 (Rambaut et al., 2018). For the Y chromosome divergence dating, using all sites resulted in poor convergence and ESS, which led us to use only SNPs that could be genotyped in all included samples (12,875 bp). We performed two independent runs, which converged to highly similar age estimates and reached ESS ≥ 323.

### *TSPY* Amplification, Sequencing and Analysis

To exclude the possibility that the discordant Y-chromosomal placement of *C. denti*, the only male representative of this species with whole genome sequencing data, was due to an aberrant sample, we amplified and sequenced a region of the Y-linked *TSPY* gene from four additional male *C. denti* individuals in two fragments, Y1 (533bp) and Y2 (445bp) (Tosi et al., 2005). Sanger sequencing was performed by the Molecular Cloning Laboratories (San Francisco, CA). Sequence chromatograms were inspected by eye and assembled using Geneious R11 11.0.5. We also extracted the corresponding regions from the mapped whole genome sequences of the *C. denti* male and one of the two *C. wolfi* males. The two regions were then concatenated, complemented with publicly available guenon *TSPY* sequences (Hart et al., 2012; Tosi et al., 2004) and aligned using MAFFT. After manual curation of the alignment, we constructed a median joining haplotype network using PopArt (Leigh and Bryant, 2015).

### Nucleotide divergence and introgression statistics

To quantify excess autosomal allele sharing between *C. denti* and *C. mitis* (indicative of gene flow), we calculated D-statistics in Dsuite/0.4 (Malinsky et al., 2021) using autosomal, biallelic SNPs. We used the rhesus macaque as outgroup, *mitis* group taxa as P3, *C. denti* as P2 and alternated between *C. mona*, *C. pogonias* and *C. wolfi* as P1. D-statistics was calculated for all possible combinations of samples, and we considered D-statistics significant if they differed from zero by more than three block-jackknife standard deviations (Z-score > 3). We used Pixy/1.2.5 (Korunes and Samuk, 2021) to calculate pairwise nucleotide divergence (*d_XY_*) between pairs of species from the *mona*, *cephus* and *mitis* species groups in non-overlapping 10 Kb windows along the genome. For *C. mitis*, we used only ssp. *opisthostictus* in these comparisons since this lineage showed the closest Y-chromosomal relationship to *C. denti*.

To search for autosomal or X-chromosomal regions that might have introgressed from *C. m. opisthostictus* into *C. denti* alongside the Y chromosome, we also calculated the *f_D_* (Martin et al., 2015) statistic in non-overlapping 10 Kb windows. The *f_D_* is related to the D-statistic but more suitable for small genomic regions with sparse data. Nevertheless, estimating introgression in short genomic regions is prone to stochasticity since there may be few informative sites. Therefore, we combined the *f_D_* statistics (setup as P1=*C. wolfi*, P2=*C. denti*, P3=*C. m. opisthostictus* and Outgroup=*M. mulatta*) with *d_XY_* to reduce the risk of false positives, and required the following criteria to be fulfilled for a window to be called as putatively introgressed from *C. mitis opisthostictus* into *C. denti*: 1) *f_D_* ≥ 95th percentile, 2) *d_XY_ C. denti* vs. *C. mitis opisthostictus* ≤ 5th percentile, 3) *d_XY_ C. denti* vs. *C. wolfi* ≥ genome wide average and 4) *d_XY_ C. wolfi* vs. *C. mitis opisthostictus* ≥ the genome wide average – 1 standard deviation. Criterion 1-3 are expected characteristics of any regions that introgressed from *C. mitis* into *C. denti*, whereas the 4th was included as a control measure to exclude regions with low absolute divergence among these lineages, since *f_D_* tends to be biased upwards in such regions (Martin et al., 2015). Any consecutive windows were merged, and genes overlapping putatively introgressed regions were identified using the Mmul_10 annotation. We used PANTHER/18.0 (Mi et al., 2021) to test for significantly overrepresented gene ontology terms among these putatively introgressed regions, using all protein coding genes of Mmul_10 as background.

### Inferences of demographic history with BPP-MSci

We used the multi-species-coalescent-with-introgression model (MSci) implemented in bpp/4.6.2 (Flouri et al., 2020) to infer divergence times, ancestral population sizes and proportion of gene flow among focal species. In this analysis, we included *C. denti*, *C. wolfi*, *C. pogonias*, *C. mona*, *C. neglectus*, *C. cephus*, *C nictitans, C. mitis opisthostictus* and *M. mulatta*, choosing the sample with the least amount of missing data in taxa with more than one available sample. We sampled a total of 1,000 loci, each 1,000 bp long, requiring them to be at least 10 Kb from the nearest gene and 50 Kb apart to avoid effects of selection and linkage, respectively. We used the topology from the ASTRAL analyses, and added three unidirectional migration bands: 1) from the ancestor of the *mitis* lineage into the ancestor of *C. pogonias*, *C. wolfi* and *C. denti*, 2) from the ancestor of the *cephus* lineage into the ancestor of *C. pogonias*, *C. wolfi* and *C. denti*, and 3) from *C. mitis* into *C. denti*. The program was set to run 20,000 iterations as burnin, and then sample every second MCMC iteration until 200,000 samples were collected. Two independent runs were performed with the same parameters, and after confirming that they converged on similar estimates, they were merged and analyzed jointly. The output tree was scaled to years and the theta values converted to effective population size (Ne), using the bppr package (Angelis and Dos Reis, 2015) assuming a mutation rate of 4.82e-9 and a generation time of 10 years (Kuderna et al., 2023). BPP assigns an Ne estimate to all branches, including those leading to hybrid nodes (e.g., the terminal branch of *C. denti* had two Ne estimates, one before and one after the incoming gene flow from *C. mitis*). To simplify our demographic model for visualization and downstream simulations, we calculated a single Ne estimate using the harmonic mean of Ne values along all branches affected by gene flow.

### Genetic simulations

To explore the probability of drift leading to the fixation of the introgressing Y chromosome, we ran coalescence simulations using msprime (Baumdicker et al., 2022). We simulated the demographic history inferred with BPP-MSCi (Figure 5), with varying amounts of gene flow from *C. mitis* into *C. denti*. For the autosomal simulations, we used the effective population size (Ne) estimates directly from the BPP output and simulated 100 loci of 1 Mb each with a recombination rate of 1e-8, always including a single pulse of 2.1 % migration from the *mitis* group ancestor into the ancestor of *C. denti*, *C. pogonias* and *C. wolfi* at 310,000 generations ago (event 1 in Figure 5), and a pulse of 8.5 % migration from the *cephus* ancestor into the same recipient population at 270,000 generations ago (event 2 in Figure 5). The third migration pulse, from *C. mitis* into *C. denti* (event 3 in Figure 5), was set to occur 100,000 generations ago (10,000 generations after the *C. denti* split from *C. wolfi*), with varying proportions from 0 to 1 %. We then used Dsuite to calculate D-statistics (P1=*C. wolfi*, P2=*C. denti*, P3=*C. mitis*, Outgroup=*M. mulatta*) and associated Z-scores. The D-statistics from simulated data were used to identify the upper limit of plausible migration proportions from *C. mitis* into *C. denti* (assuming that gene flow was a neutral process), given our empirical estimates.

We also simulated a non-recombining 10 Kb locus to mimic the Y chromosome under the same demographic history as above, dividing the effective population sizes by four since the ratio Ne_Ychrom_ / Ne_Autosomes_ = 0.25 (Laporte and Charlesworth, 2002).

Considering that guenon males disperse more than females, it is likely that gene flow is predominantly driven through male migration. Under the conservative assumption that only males migrate, the effective Y-chromosomal migration rate would be twice that of the autosomes. Therefore, we doubled the migration proportions from *C. mitis* into *C. denti* compared to the autosomal simulations, to make them directly comparable. We simulated 100 replicates of 1,000 simulations per migration rate (0-2 % with a stepwise increase of 0.1 %), and counted the number of times *C. denti* and *C. mitis* formed a monophyletic clade.

To test how selection affects the probability of Y chromosome fixation in *C. denti*, we ran forward simulations in SLiM (Haller and Messer, 2019). We simulated a single population of 200,000 individuals for 100,000 generations (mimicking the *C. denti* lineage after the split from *C. wolfi*), and used the built-in functionality of modeling Y chromosome to track the frequency of a novel Y-linked allele introduced in the first generation. We tested selection coefficients (s) ranging from 0-0.01, and initial Y chromosome frequencies (equivalent to incoming male migration) in the range of 0.1-1 % (range informed by previous neutral simulations, Figure 6A). We ran 100 replicate simulations for each frequency and selection coefficient, counting the number of times the allele was lost, fixed or still segregating after 100,000 generations.

### Identifying candidate genes under selection and possible drive elements

We used several approaches to explore putative genes under positive selection that might have driven the introgressing Y chromosome to fixation in *C. denti*. First, we identified protein coding genes annotated on the rhesus macaque Y chromosome which showed differentially fixed amino acid sequence in *C. mitis opisthostictus* and *C. denti* compared to *C. wolfi* and *C. pogonias*. We used custom Python scripts to translate the transcripts from each gene into amino acid sequences and to count the number of fixed differences for each gene, after excluding genes with internal stop codons in focal lineages. Second, we tested whether a model of adaptive or neutral evolution along the *C. denti* and *C. mitis* branch was a better fit for genes with fixed amino acid differences, using codeml in paml/4.9j and HyPhy/2.5.51 (Kosakovsky Pond et al., 2019). Codeml was run using the branch-site model: A model allowing for positive selection on a subset of sites along a specified set of foreground branches was compared to a model of neutral evolution. We tested the following foreground branches, based on the Y-chromosomal topology: 1) the ancestral *mitis* group branch and all descendants (including *C. denti*); 2) the ancestral *mitis* group branch, the ancestral *C. mitis* + *C. denti* branch and the terminal branches of *C. mitis* and *C. denti*; 3) the ancestral *C. mitis* + *C. denti* branch and their respective terminal branches; and 4) only the terminal branch of *C. denti*. For these analyses, we used a single individual for each *Cercopithecus* species in our data set and *Chlorocebus sabaeus* as the outgroup, choosing the sample with the least amount of missing data. HyPhy was run using the ‘meme’ algorithm (Murell et al. 2012), on a concatenated alignment of the Y chromosome genes using the same species.

We also identified sex-chromosomal regions with increased coverage, indicative of copy-number expansions, as putative candidates for meiotic drive elements (Baird et al., 2023; Hughes et al., 2020). To this end, we calculated the average mapping depth in sliding windows of 5 Kb (step size 1 Kb) along the rhesus macaque X and Y chromosomes. The coverage was normalized for each sample and chromosome by dividing the window depths by the chromosome-wide average. Next, we calculated the coverage ratio of *C. denti* to *C. wolfi*, alternating through all pairwise comparisons of individuals between these species. If meiotic drive through a copy number expansion of specific Y-linked genes occurred in *C. denti*, such regions are expected to show higher coverage in *C. denti* genomes relative to its sister *C. wolfi*. In this case, we also expect to observe compensatory increase in copy number on the X-chromosome.

## Acknowledgements

We thank Stuart JE Baird for useful discussions on mammalian Y chromosome introgression, Terese Hart for logistical support for sample collection in the Democratic Republic of Congo, and Louis Rugyerinyange, Felix Mulindahabi and Beth Kaplin for logistical support for sample collection at Nyungwe National Park. We acknowledge Faustin Kahindo, Maurice Emetshu, Gilbert Paluku, Peter-Philip Niehoff and James Gray for assistance with sample collection in the field. We thank Stephen Nash for providing the guenon illustrations in Figure 1 and 2. Sequencing was performed by the SNP&SEQ Technology Platform in Uppsala. The facility is part of the National Genomics Infrastructure (NGI) Sweden and Science for Life Laboratory. The SNP&SEQ Platform is also supported by the Swedish Research Council and the Knut and Alice Wallenberg Foundation. The computations were enabled by resources in projects SNIC 2022/6-325 and SNIC 2022/5-561, provided by the Swedish National Infrastructure for Computing (SNIC) at Uppsala University (UPPMAX), partially funded by the Swedish Research Council through grant agreement no. 2018-05973. The project was supported by the Swedish Research Council VR (2020-03398) grant to K.G., Zoologiska Stiftelse grants to A.J., Margot Marsh Biodiversity Foundation and FAU Foundation, Inc. to K.M.D. A.T. acknowledges funding from NSF award #1718715.

## Data availability

The sequence data generated for this project are available at ENA/SRA under accession number XXX. Accession numbers of downloaded publicly available sequence data are listed in Table S1. Scripts are available at https://github.com/axeljen/denti_ychrom_scripts.

