## Supplementary figures for "Breaking the rule: An exceptional Y chromosome introgression between deeply divergent primate species"

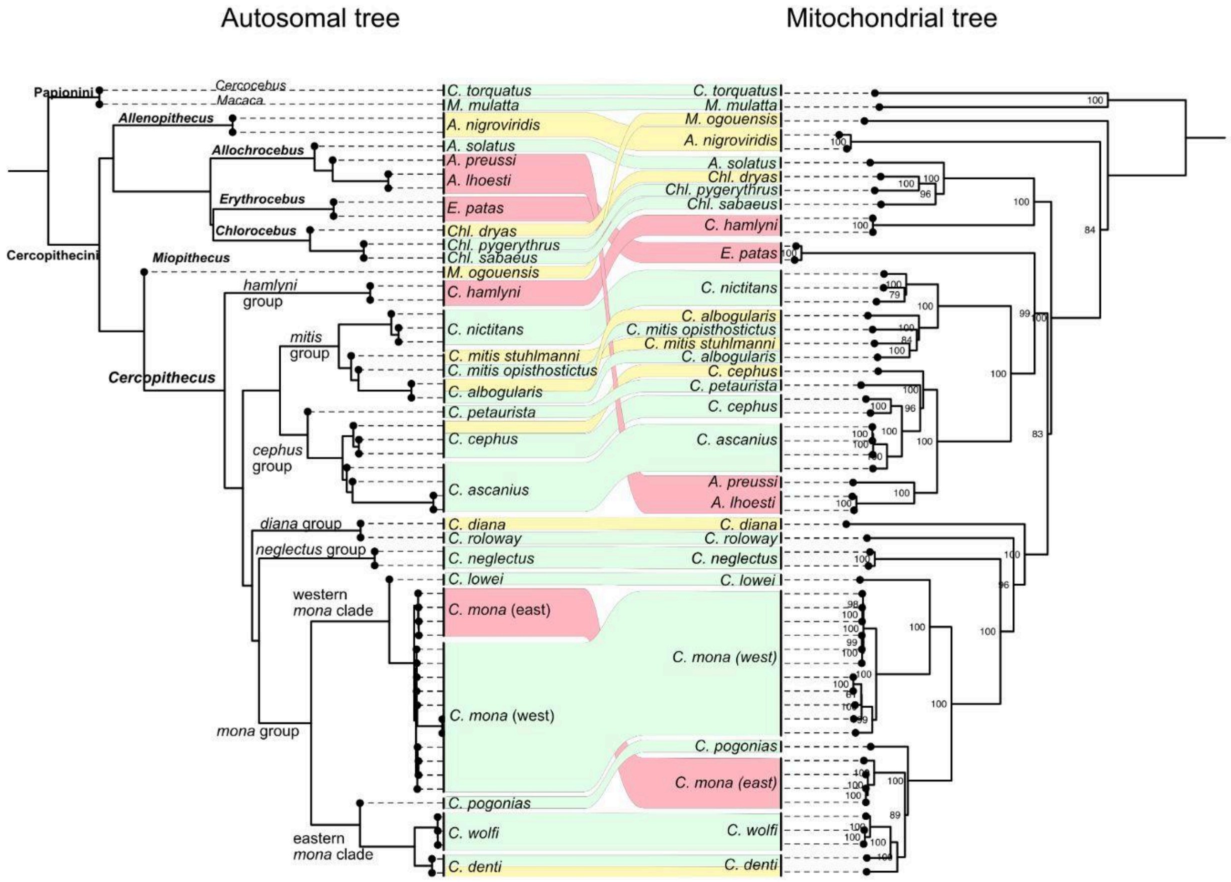


**Figure S1. Mito-nuclear conflicts.** Concordant phylogenetic positions between the autosomal and the mtDNA trees are shown by green connections. Shallow discordances that can be the result of both ILS and introgression are shown in yellow, whereas deep discordances that cannot be explained by ILS and require introgression are shown in salmon.

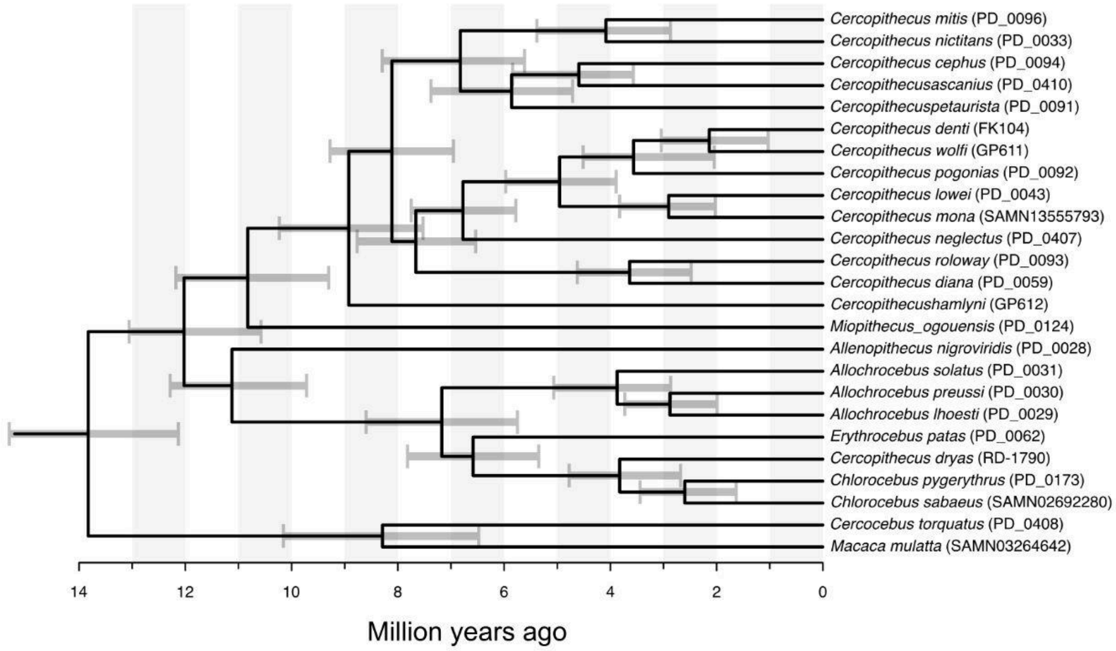


**Figure S2. Autosomal divergence date estimates using MCMCTree.** Divergence dates were estimated using a single sample per species (ID in parenthesis, table S1). Gray bars correspond to 95% HPD credible intervals.

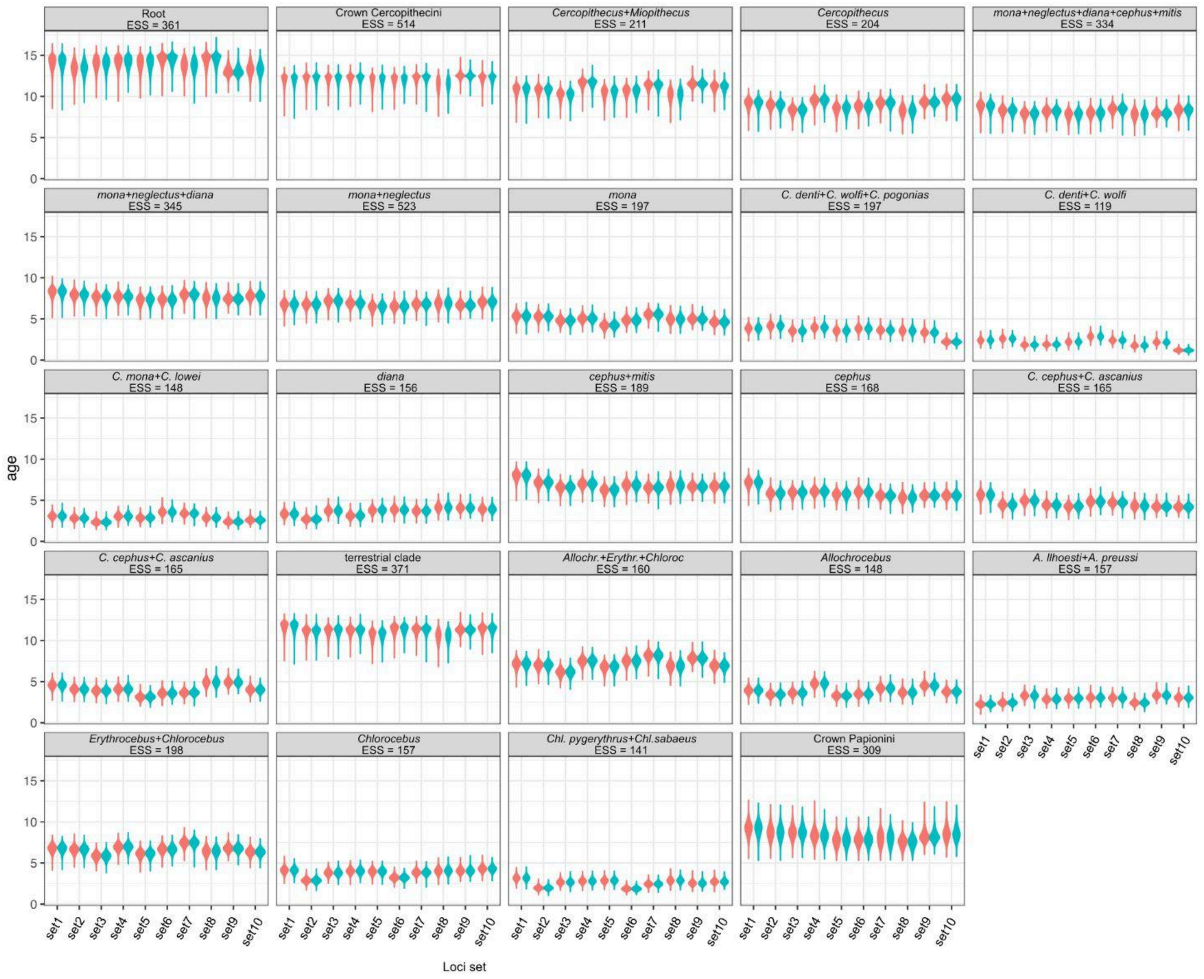


**Figure S3. Autosomal date estimate distribution/MCMCTree convergence.** Age distributions among all autosomal MCMCTree runs shown as violin plots. The analysis was run on ten independent loci (set 1-10), and each set was analyzed in two independent runs (the two runs for each locus are color coded red and green). Age on the Y-axis given in million years. The nodes to which the age estimates apply are given above each panel, together with the obtained Effective Sample Size (ESS).

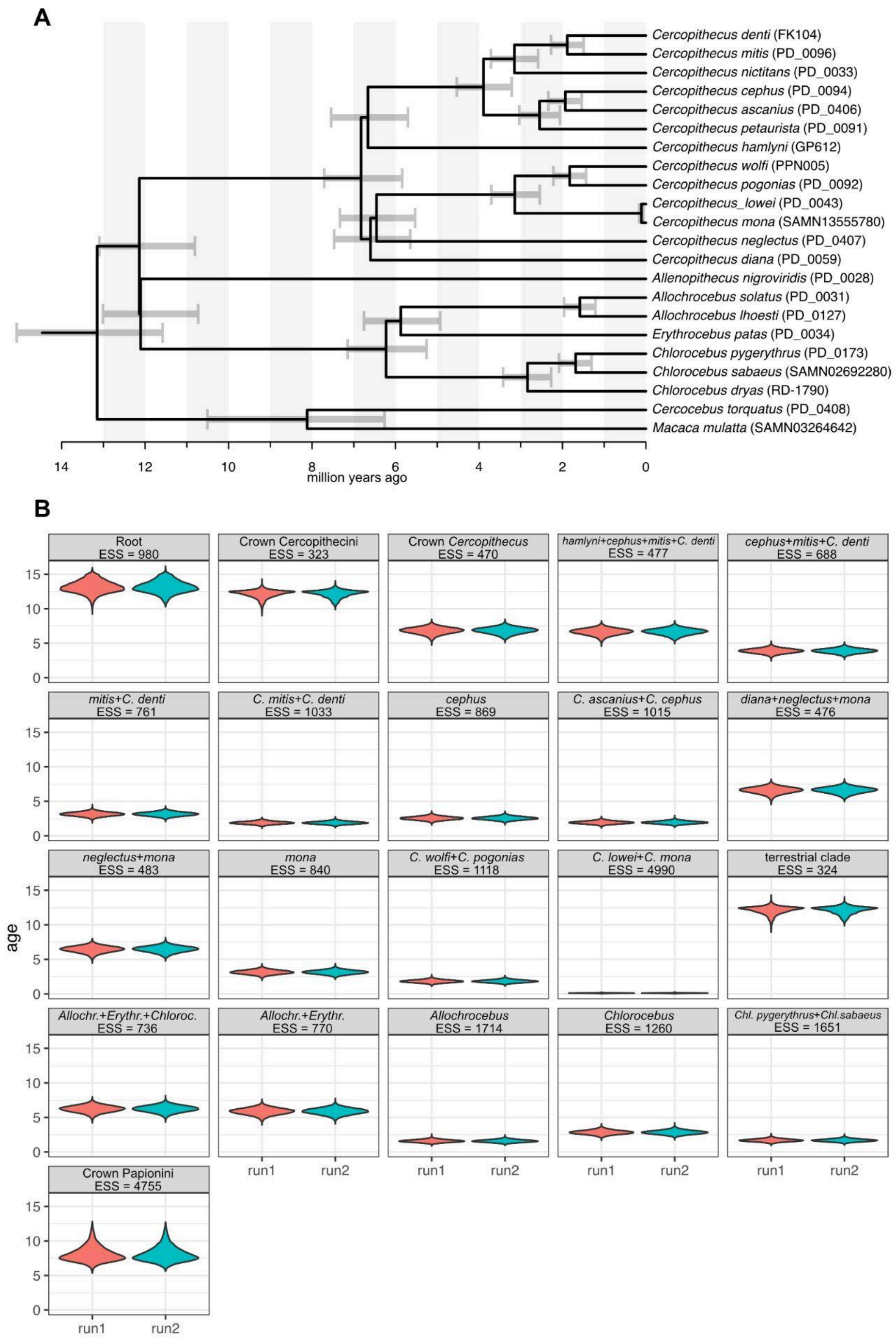


**Figure S4. Y-chromosomal date estimate distribution/MCMCTree convergence.** Age distributions among all Y-chromosomal MCMCTree runs shown as violin plots. The analysis was run on a Y chromosome alignment (see Methods), analyzed in two independent runs. Age on the Y-axis given in million years. The nodes to which the age estimates apply are given above each panel, together with the obtained Effective Sample Size (ESS).

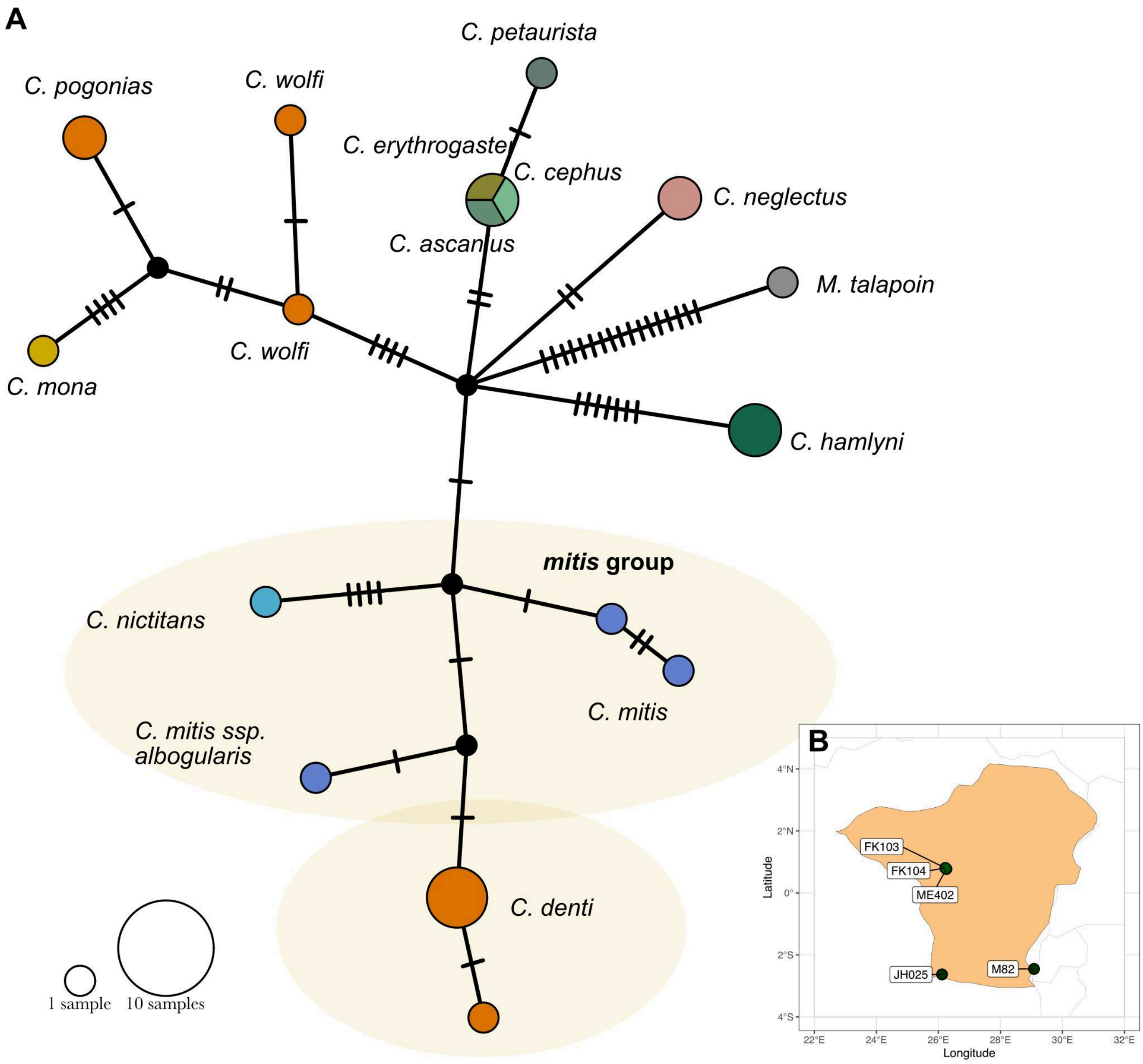


**Figure S5. Median joining network based on 898 bp from the Y-linked TSPY locus.** The network includes five *C. denti* individuals, sampled across the distribution range, and shows that the introgressed *mitis*-like Y-chromosome is present in all study samples. Inset map shows the distribution range of *C. denti* (orange) with points showing the sampling locations of the sequenced individuals.

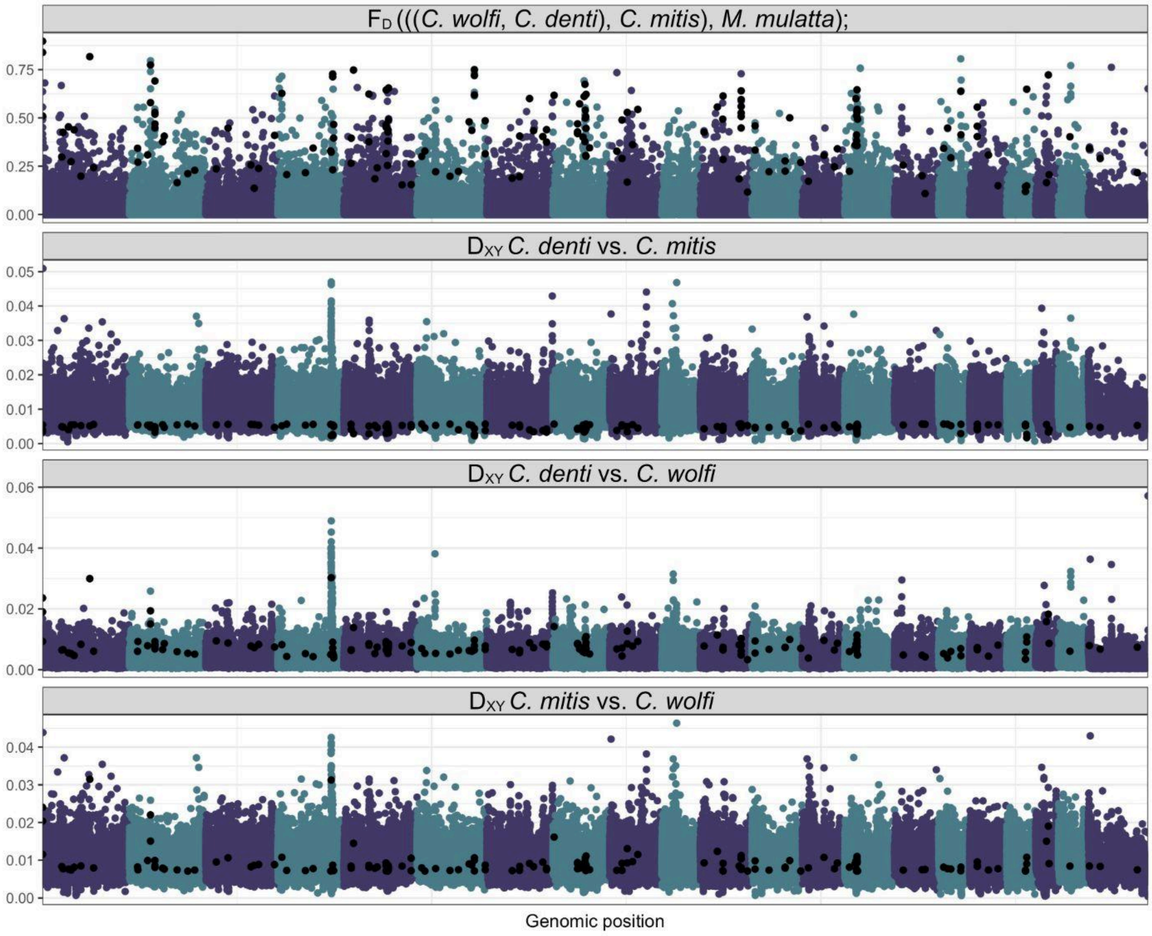


**Figure S6. Genomic landscape of introgression from *Cercopithecus mitis opisthostictus* into *C. denti*.** Top panel shows $F_D$(P1=*C. wolfi*, P2=*C. denti*, P3=*C. mitis*, outgroup=*M. mulatta*), and the remaining panels show relevant $D_{XY}$ comparisons. *C. m. opisthostictus* was used as the *C. mitis* representative in all comparisons. Windows called as putatively introgressed are plotted as black dots across all panels, and were called if they satisfy all of the following criteria: 1) $F_D$ > 95th percentile, 2) $D_{XY}$ *denti-mitis* < 5th percentile, 3) $D_{XY}$ *denti-wolfi* > genome wide average and 4) $D_{XY}$ *wolfi-mitis* > genome wide average - 1 sd. Alternating colors depict chromosomes in the reference *M. mulatta* genome.

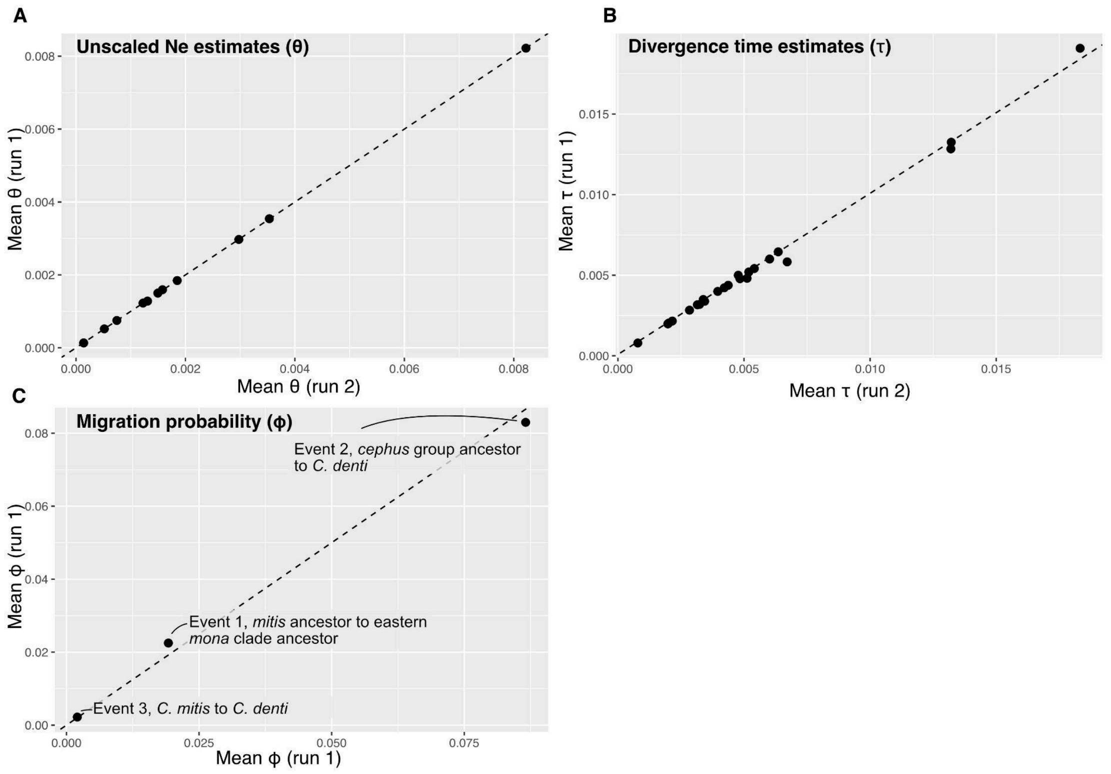


**Figure S7. BPP convergence between two independent runs of the MSCi model.** Comparisons between two independent BPP-MSci runs (Figure 5 in the main text). A) Theta (θ) estimates, where each point represents the estimated unscaled effective population size (Ne) for one branch in the tree. B) Unscaled divergence time estimates, tau (τ), the estimated divergence time for each internal node is represented by one point. C) Migration probability estimates, phi (φ), for the three modeled gene flow events (event 1, 2 and 3 in Figure 5 in the main text).

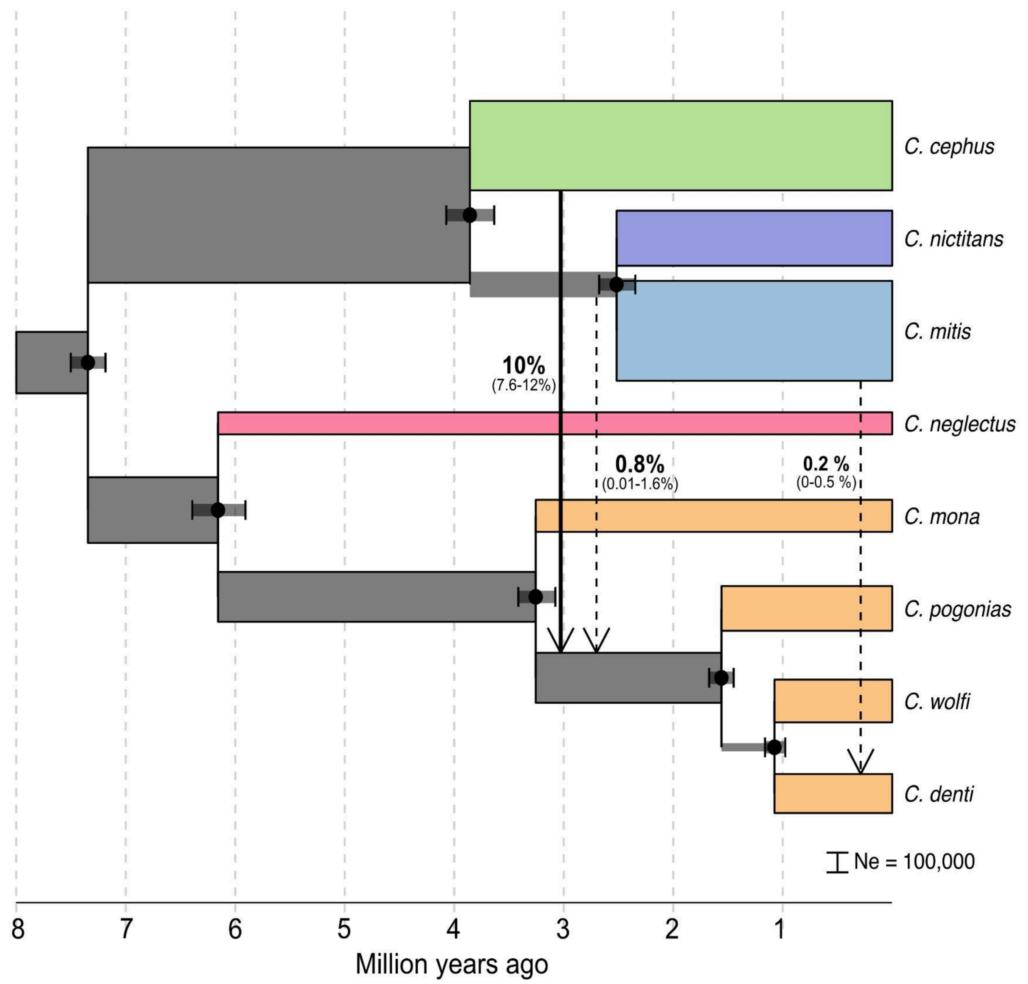


**Figure S8.** Demographic history of focal lineages using BPP and the same model as in main Figure 5, except that the gene flow event from *C. cephus* into the *C. pogonias/wolfi/denti* ancestor precedes that from the *mitis* group ancestor. Arrows show the modeled gene flow events, labelled with the estimated migration proportion and 95 % highest posterior distribution credible interval. Branch lengths and widths were scaled to years and effective population size, respectively, using a generation time of 10 years and a mutation rate of 4.82e-9.

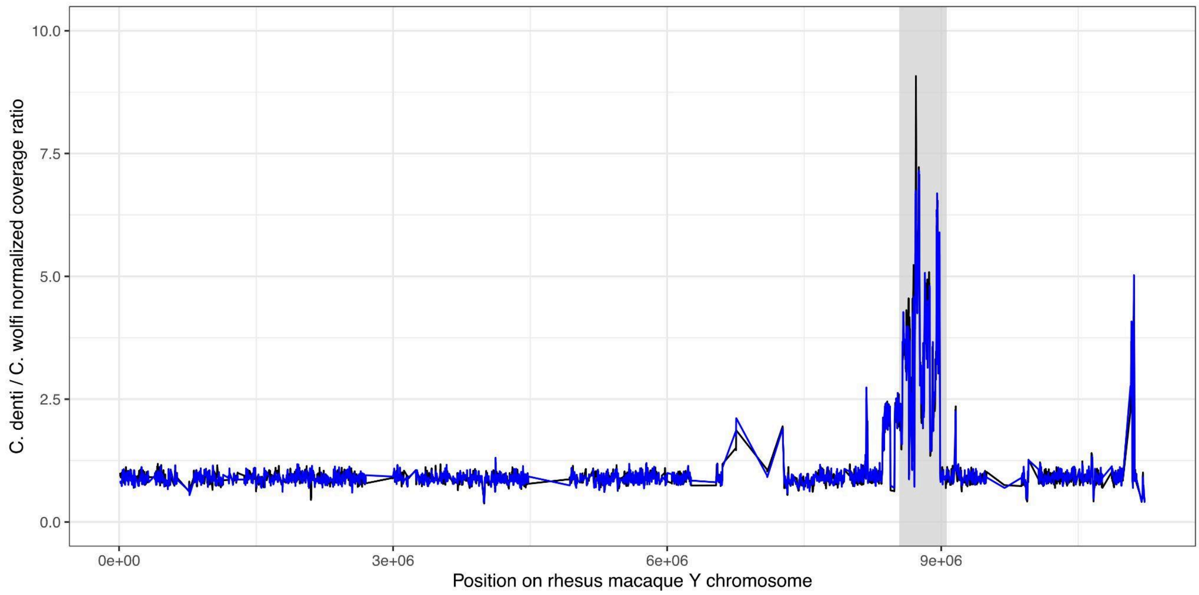


**Figure S9**. Normalized coverage ratio *C. denti* / *C. wolfi* along the rhesus macaque Y-chromosome, calculated separately for both included *C. wolfi* males (blue and black line, respectively). The grey rectangle highlights the region of increased coverage in *C. denti* compared to *C. wolfi*.

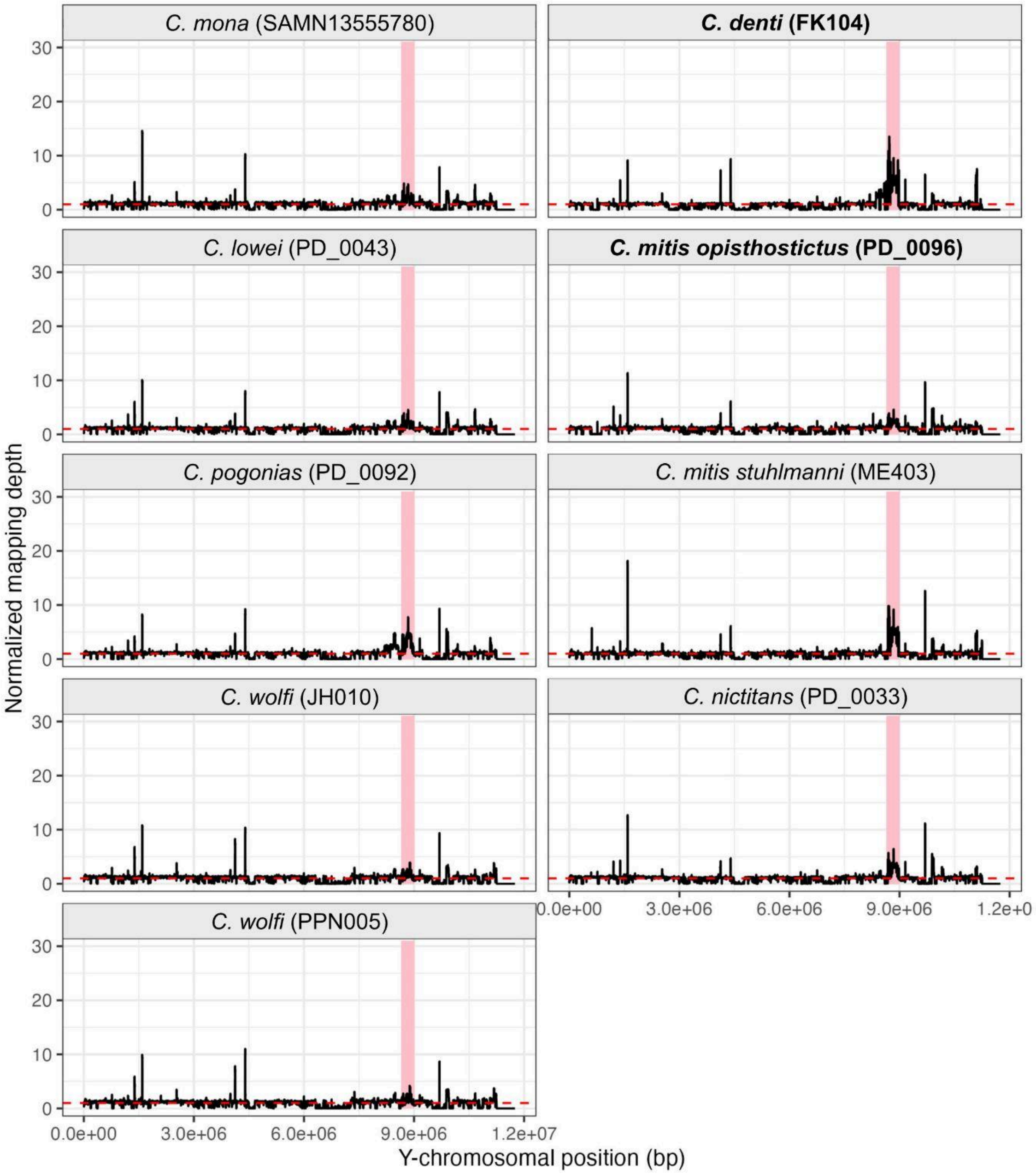


**Figure S10.** Mapping coverage along the macaque Y-chromosome, for a subset of guenon species, normalized by average Y-chromosomal coverage. Left column shows the Y-coverage among *mona* group lineages that retained the ancestral allele, and the right column shows the coverage in *C. denti* and *mitis* group taxa. Horizontal dashed line shows the normalized coverage value of one. The region showing high variation in Y-chromosomal coverage is highlighted by red rectangles.

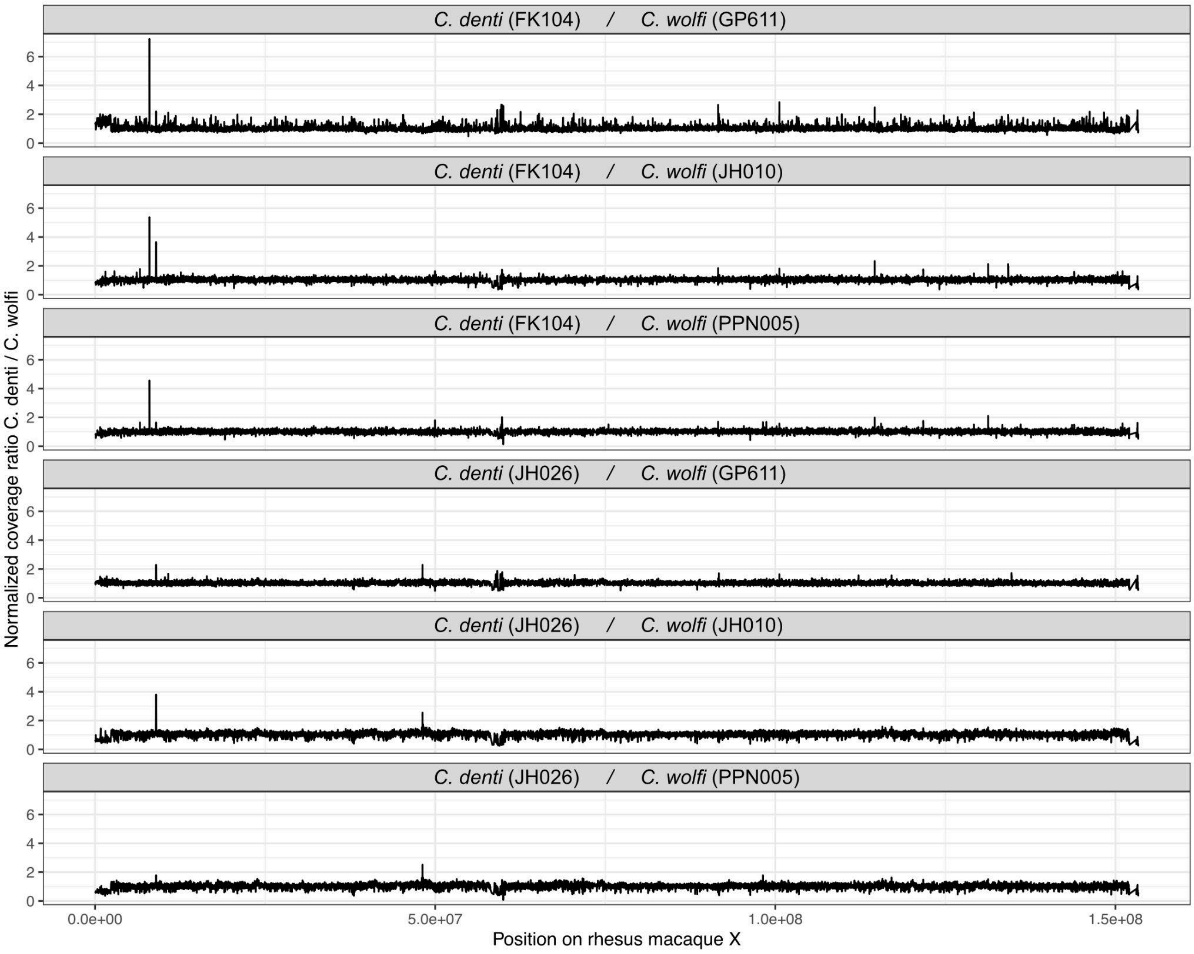


**Figure S11**. Normalized coverage ratio between all combinations of *C. denti* and *C. wolfi* samples in windows along the rhesus macaque X chromosome. Coverage was normalized by dividing the mean coverage in each window with the X-chromosome wide average for each sample.

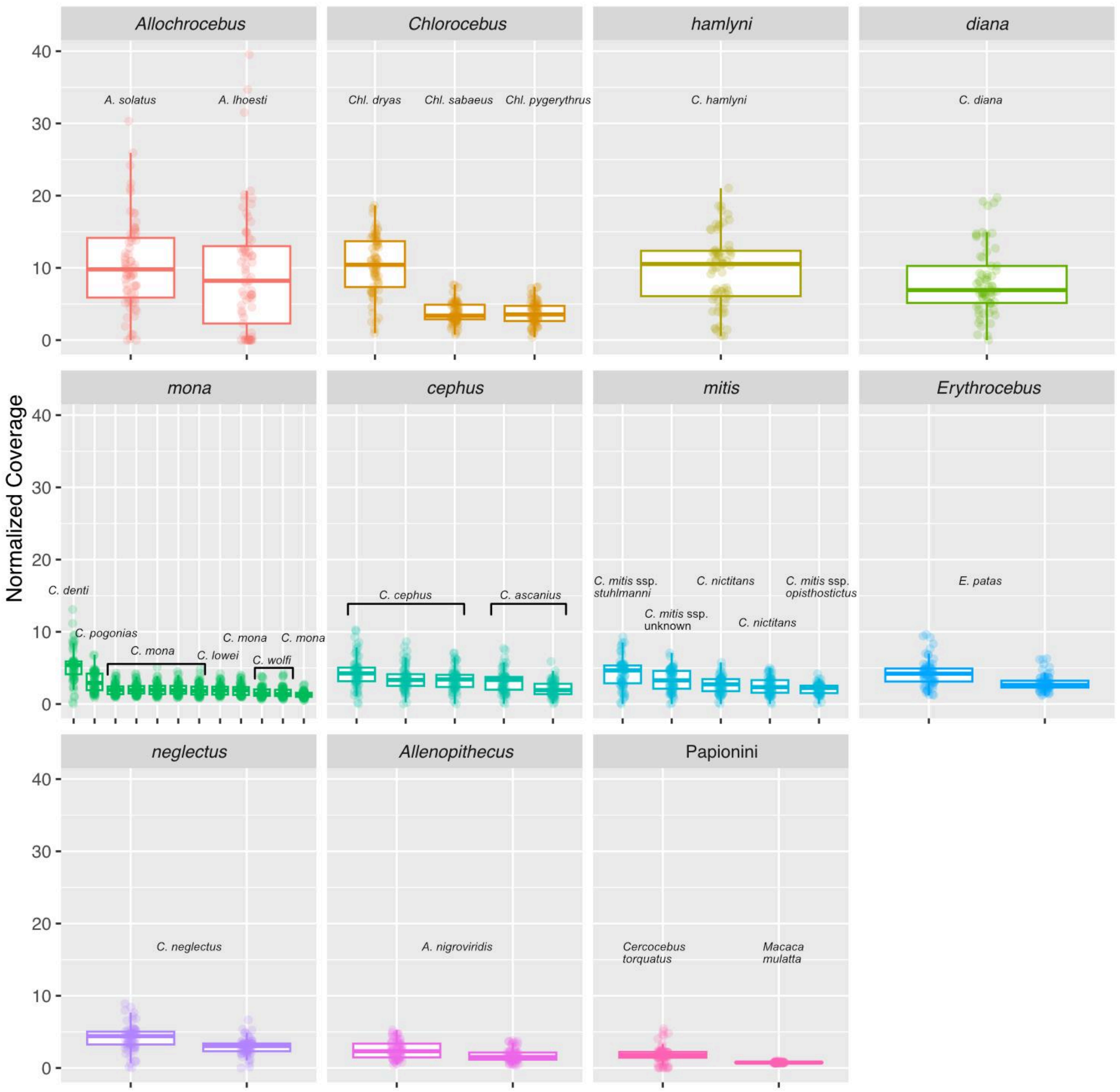


**Figure S12.** Mapped read depth in the putative ampliconic region on the guenon Y-chromosome, normalized by the average Y-chromosomal coverage. Each point represents a non-overlapping 5 kb window in the rhesus macaque Y-chromosomal region 8,650,000-9,000,000 bp.
